# Impact of Δ^9^-Tetrahydrocannabinol and oxycodone co-administration on measures of antinociception, dependence, circadian activity, and reward in mice

**DOI:** 10.1101/2023.12.04.569809

**Authors:** Richard A. Slivicki, Justin G. Wang, Vy Trinh Tran Nhat, Alexxai V. Kravitz, Meaghan C. Creed, Robert W. Gereau

**Affiliations:** Washington University Pain Center and Department of Anesthesiology, Washington University, St. Louis, MO; Neuroscience Graduate Program, Division of Biology & Biomedical Sciences, Washington University, St. Louis, MO; Department of Psychiatry, Washington University, St. Louis, MO; Department of Neuroscience, Washington University, St. Louis, MO; Department of Biomedical Engineering, Washington University, St. Louis, MO

## Abstract

Oxycodone is commonly prescribed for moderate to severe pain disorders. While efficacious, long-term use can result in tolerance, physical dependence, and the development of opioid use disorder. Cannabis and its derivatives such as Δ^9^-Tetrahydrocannabinol (Δ^9^-THC) have been reported to enhance oxycodone analgesia in animal models and in humans. However, it remains unclear if Δ^9^-THC may facilitate unwanted aspects of oxycodone intake, such as tolerance, dependence, and reward at analgesic doses. This study sought to evaluate the impact of co-administration of Δ^9^-THC and oxycodone across behavioral measures related to antinociception, dependence, circadian activity, and reward in both male and female mice. Oxycodone and Δ^9^-THC produced dose-dependent antinociceptive effects in the hotplate assay that were similar between sexes. Repeated treatment (twice daily for 5 days) resulted in antinociceptive tolerance. Combination treatment of oxycodone and Δ^9^-THC produced a greater antinociceptive effect than either administered alone, and delayed the development of antinociceptive tolerance. Repeated treatment with oxycodone produced physical dependence and alterations in circadian activity, neither of which were exacerbated by co-treatment with Δ^9^-THC. Combination treatment of oxycodone and Δ^9^-THC produced CPP when co-administered at doses that did not produce preference when administered alone. These data indicate that Δ^9^-THC may facilitate oxycodone-induced antinociception without augmenting certain unwanted features of opioid intake (e.g. dependence, circadian rhythm alterations). However, our findings also indicate that Δ^9^-THC may facilitate rewarding properties of oxycodone at therapeutically relevant doses which warrant consideration when evaluating this combination for its potential therapeutic utility.

## Introduction

Oxycodone remains one of the most prescribed analgesics for moderate to severe pain disorders(Dowell et al., 2022). Oxycodone, like other opioids, exerts its analgesic actions primarily through the µ-opioid receptor(Beardsley et al., 2004; Olkkola et al., 2013). While effective in the short term, long-term use of µ-opioid agonists can result in a myriad of unwanted side-effects which include tolerance, changes in the sleep-wake cycle, physical dependence, and the development of opioid use disorder(Strang et al., 2020). Thus, there is an urgent need to develop alternative therapeutic interventions for pain.

Cannabis and its derivatives such as Δ^9^-Tetrahydrocannabinol (Δ^9^-THC) have been suggested as a potential therapeutic for pain. Cannabis and cannabis-derivative containing products have been recently made available for medicinal use in most states within the United States(Haroutounian et al., 2021). In a recent survey among adults with chronic pain who reside in in states with legalized medicinal cannabis, 30% of individuals reported using cannabis to treat their pain(Bicket et al., 2023). In preclinical studies, Δ^9^-THC administration produces antinociception and exerts efficacy in evoked and non-evoked pain behaviors across several different pain models(Abraham et al., 2020; Soliman et al., 2021). Δ^9^-THC exerts these actions primarily through cannabinoid type-1 (CB1) and type-2 (CB2) receptors(Woodhams et al., 2017). However, like µ-opioid agonists, prolonged administration of Δ^9^-THC and other CB1 agonists results in antinociceptive tolerance and dependence thereby limiting the therapeutic window of CB1 agonists(Ramaekers et al., 2020). Better approaches to maximizing the therapeutic potential of cannabinoid-based therapies are therefore needed.

There is considerable interest in combination therapies of cannabinoids and opioids for analgesia. CB1 and µ-opioid receptors are co-expressed in areas throughout the pain neuraxis where they are poised interact in the context of analgesia(Wilson-Poe et al., 2021; Yu et al., 2022) and analgesic tolerance(Wilson-Poe et al., 2012; Wilson-Poe et al., 2013). Indeed, a wealth of preclinical evidence generally suggests co-administration of CB1 and µ-agonists result in enhanced antinociception, and synergy across different pain models(Welch and Stevens, 1992; Cichewicz et al., 1999; Cichewicz and McCarthy, 2003; Cichewicz et al., 2005; Wilkerson et al., 2016; Wilkerson et al., 2017; Maguire and France, 2018; Slivicki et al., 2018; Slivicki et al., 2020; Nielsen et al., 2022; Toniolo et al., 2022; Yu et al., 2022). A recent meta-analysis reported 3.5-fold reduction in the morphine dose required to produce analgesia when combined with Δ^9^-THC in rodent studies(Nielsen et al., 2022). This facilitation of antinociception is also observed in nonhuman primates(Maguire et al., 2013; Maguire and France, 2014; Maguire and France, 2018; Gerak et al., 2019; Nilges et al., 2020). In humans, combined treatment with oral oxycodone and smoked cannabis resulted in enhanced antinociception in the cold pressor test relative to either drug in isolation(Cooper et al., 2018). A recent review concluded the clinical evidence of cannabinoid and opioid combinations are largely mixed in their analgesic effectiveness(Babalonis and Walsh, 2020). Studies providing evidence for or against the use of specific cannabinoid/opioid combinations in a therapeutic context are therefore likely to be highly useful.

In addition to pain, there is evidence that cannabis legalization has an impact on opioid misuse and overdose. Studies indicate that there are fewer opioid overdose deaths in states where recreational cannabis is legal(Shi, 2017; Marinello and Powell, 2023). However, other reports suggest that access to cannabis may increase the risk of opioid misuse(Ali et al., 2023; Rhew et al., 2023) potentially resulting in the development of opioid use disorder. In addition to pain-relevant areas, CB1 and µ-opioid receptors are co-expressed in areas such as the ventral tegmental area and nucleus accumbens whose activity is critical for the rewarding properties of reinforcing drugs(Wenzel and Cheer, 2018). Indeed, Δ^9^-THC has been reported to both facilitate(Solinas et al., 2004) and reduce(Gutierrez et al., 2022) heroin self-administration. Recently, it was demonstrated that injected or vaporized Δ^9^-THC reduced oxycodone self-administration(Nguyen et al., 2019). Similarly, in non-human primates it has been reported that Δ^9^-THC administration decreases heroin responsivity(Maguire and France, 2016), but has no effect on fentanyl reward(Carey et al., 2023) suggesting a potential ligand-specific interaction. In humans studies, smoked cannabis resulted in a facilitation of drug liking for oxycodone(Cooper et al., 2018), however oral administration of the Δ^9^-THC-conjugate dronabinol reduced drug liking for oxycodone(Lofwall et al., 2016). The two studies differ in their route of administration and dose of oxycodone, further suggesting these interactions are complex and highly dependent on dose and route of administration. Thus, there are clear interactions between Δ^9^-THC and opioids in the context of reward that require consideration when evaluating potential therapeutic implementation. CB1 and µ-opioid receptors are also co-expressed in areas relevant to drug withdrawal(Scavone et al., 2013). CB1 agonists, allosteric modulators and endocannabinoid degradation inhibitors reduce measures of µ-opioid dependence and withdrawal(Lichtman et al., 2001; Schlosburg et al., 2009; Ramesh et al., 2011; Wilkerson et al., 2017; Dodu et al., 2022). In humans, the Δ^9^-THC conjugate dronabinol has been shown to suppress opioid withdrawal signs in opioid-dependent individuals(Bisaga et al., 2015). However, in another study dronabinol exhibited unwanted effects such as a “heart racing” feeling and tachycardia at doses effective for reducing withdrawal measures in opioid-dependent individuals(Lofwall et al., 2016). Thus, experiments evaluating how co-treatment of oxycodone and Δ^9^-THC interact in the context of drug withdrawal are likely to be translationally informative.

Prior literature indicates that cannabinoids and opioids functionally interact to affect a wide array of different outcome measures. Ligand, dose range and dosing regimen can all influence such interactions. Based on these studies, we set out to evaluate the impact of co-administration of oxycodone with Δ^9^-THC in a variety of behaviors related to antinociception, tolerance, dependence, circadian activity, and reward.

## Methods

### Subjects

All experiments used C57BL/6J male and female mice bred in-house or purchased from Jackson laboratories (Bar Harbor, ME). Mice were 8 to 10 weeks of age at the start of experiments. Animals were group housed 3-5 per cage in all experiments aside from the hotplate combination dosing experiments. Animals were maintained in a temperature-controlled facility with a 12-hour light–dark cycle (lights on at 06:00-18:00 hours) and given ad libitum food and water. All experimental procedures were approved by the Washington University Institutional Animal Care and Use Committee and followed the guidelines of the International Association for the Study of Pain. Mice were randomly assigned to experimental conditions. Male and female cohorts were always tested separately. Behavioral testing occurred between 8 AM and 6 PM (2:00 and 12:00 ZT).

### Drugs and chemicals

Δ^9^-THC (NIDA drug supply program) was provided in ethanol (200 mg/ml), and dissolved in a vehicle of 4.75% ethanol, 5% Koliphor EL, and 90% sterile saline and administered at volume of 5 ml/kg. For this vehicle, a stock of Koliphor and saline was separately mixed and then added to the corresponding amount of ethanol to yield the final injectable vehicle. Oxycodone (Sigma-Aldrich) was dissolved in sterile saline and administered at a volume of 5 ml/kg. For experiments which involved combinations of Δ^9^-THC and oxycodone both compounds were dissolved in the same vehicle (4.75% ethanol, 5% Koliphor EL and 90% sterile saline) at 2.5 ml/kg. Equal parts of each solution were combined to generate Δ^9^-THC and oxycodone cocktail which was injected at a volume of 5 ml/kg.

### Hotplate test of antinociception

The hotplate test was used to evaluate antinociception as both oxycodone and Δ^9^-THC have been demonstrated to be antinociceptive in this assay(Zhang et al., 2016; Alegre-Zurano et al., 2020). In brief, mice were placed on a hot plate (Model PE34 Series 8, IITC Life Science Inc.) set at 55°C. The latency to nocifensive response by any paw was recorded. A maximal cutoff latency of 15s seconds was used to prevent tissue damage.

### Measuring oxycodone and Δ^9^-THC acute antinociception and tolerance

Oxycodone or Δ^9^-THC was administered at doses of 1, 3, 10 or 30 mg/kg twice daily during the light cycle at approximately 0830 and 1730 for 5 consecutive days. Hotplate thresholds were measured 30, 60, 90, 180, 360 and 1440 mins after the first daily injection on days 1, 3 and 5 of repeated dosing. Dose-response curves were generated from the 30 min timepoint on day 1 of dosing. Groups received oxycodone (3 mg/kg s.c.), Δ^9^-THC (3 mg/kg s.c.), combination of oxycodone (3 mg/kg s.c.) and Δ^9^-THC (3 mg/kg s.c.), or vehicle. Drug treatment and hotplate testing were conducted as described above.

### Evaluation of µ-opioid receptor dependence

Following repeated dosing with the assigned condition, mice were treated with a final dose of the assigned treatment condition followed by naloxone (3 mg/kg s.c.) 1 hr later. Immediately following naloxone injection, mice were placed into a plexiglass observation chamber and were recorded with a Sony Handicam set at 30 frames per second with 1080p resolution. DeepLabCut^TM^ (DLC, version 2.2b6[46,47]) was used to conduct markerless pose estimation as described in(Slivicki et al., 2023). The network was built from 20 frames extracted from 30 videos using kmeans clustering and labelled with the following 9 body parts: left ear, right ear, left forepaw, right forepaw, left hind paw, right hind paw, snout, tail base, back. The training fraction was set to 0.95, and the resnet_50 network was trained for 800,000 iterations. A train error of 1.82 and test error of 6.67 were achieved with a cutoff value of p=0.6. From this point the trained networked was locked and used to analyze subsequent videos.

Simple behavioral analysis (SimBA)(Nilsson et al., 2020) was used to process DLC data as previously described(Slivicki et al., 2023). Videos were cropped for occasions which contained frames of jumping and other behaviors, typically around a minute long. Frames were labelled for jumping behavior (any instance in which all 4 paws left the platform). Random forest classification was then used to generate a behavioral classifier for jumping behavior with the probability set to 0.9 and behavior bout 35 ms. This classifier was then applied to subsequent videos to generate the total number of jumping instances.

### Home cage activity monitoring using passive infrared (PIR)

To evaluate circadian activity, mice were single housed and a PIR-based activity sensor (Pallidus MR1, Saint Louis MO) was placed in the homecage to detect changes in activity as reported previously(Slivicki et al., 2023). Briefly, the PIR sensor tested for activity in the homecage 5 times per second and data were aggregated as the % of time that the mouse was active in each minute. To evaluate the acute effect of a single drug injection on activity, % of time active was analyzed as 5 min bins (Supplemental Figure 1). To evaluate general effects on circadian rhythm throughout each experimental phase, activity % was analyzed as 1 hr bins throughout the entire light:dark cycle (See Supplemental Figure 2 for collapsed averages throughout each phase of the experiment). Data were processed on the MR1 and transmitted wirelessly to a cloud-based server for storage and visualization. Animals were monitored continuously for 2 days prior to the dosing period. During the dosing period animals were injected with vehicle, oxycodone (3 mg/kg s.c.), Δ^9^-THC (3 mg/kg s.c.) or a combination of oxycodone (3 mg/kg s.c.) and Δ^9^-THC (3 mg/kg s.c.) twice daily during the light cycle as detailed above.

### Conditioned place preference (CPP) and locomotor sensitization

CPP was performed as described(Land et al., 2008). The apparatus consisted of two chambers with distinct visual cues (vertical or horizontal black and white stripes) and one center chamber. Each chamber was filled with bedding, and the chambers were wiped down with water between sessions. **Habituation and baseline chamber preference:** On days 1 and 2, animals were allowed access to all three chambers for 30 min and the baseline values for time spent in each chamber were derived from day 2. Vehicle/drug chamber pairings for each mouse were then randomly assigned. **Pairing:** Days 3-5 served as chamber pairing days. On these days animals were first injected with vehicle in the AM and confined to the designated chamber for 30 min. At least 4 hrs later animals were injected with the assigned drug conditioned and were confined to the opposite chamber for 30 min. **Post-test:** On day 6, animals were again allowed free access to all three chambers for 30 min.

Video was recorded from the top of each chamber at rate of 25 fps using Bonsai open-source software(Lopes et al., 2015). Videos were then analyzed using Ethovision 14 (Noldus Information Technology). Ethovision was used to calculate the amount of time spent in each chamber and the amount of distance travelled during conditioned sessions.

### Statistical analysis

The experimenters (RS, JW) were blind to behavioral treatment condition. All data were analyzed using Graphpad Prism 8.0 and Excel. Raw data for hotplate analysis and ED50 calculations were converted to % baseline responding (i.e. prior to CFA treatment) using the following formula: (Experimental Value – Hotplate Baseline)/(15 – Hotplate Baseline). ED50 values were generated using nonlinear regression analysis in GraphPad 9.0. 5 min bins of activity counts were used to demonstrate acute changes in activity (supplemental figure 1). For circadian experiments, activity counts were comprised of 1 hr bins for the entirety of the light:dark cycle. 24 hr data was transformed to area under the curve (AUC) for each light:dark period (12 hrs). Total percentage of time active during the light phase % was calculated by dividing the AUC of the light activity during the 12 hour ‘light-on’ period by the total AUC for that day. Behavioral data were analyzed via an ANOVA (two-way or one-way) followed by Tukey’s or Sidak’s post-hoc tests. Relevant statistically significant comparisons are reported in the text, represented in the figure legends. Detailed statistics are reported in the supplemental tables.

## Results

### Oxycodone and Δ^9^-THC produce dose-dependent antinociception in the hotplate test

To evaluate antinociceptive efficacy and tolerance, separate groups of male and female mice were administered oxycodone (0,1,3,10, and 30 mg/kg s.c.), Δ^9^-THC (0,3,10,30 mg/kg s.c.) or their respective vehicles twice daily for a period of 5 days (see Figures 1A, 2A for timeline).

**Figure 1.**
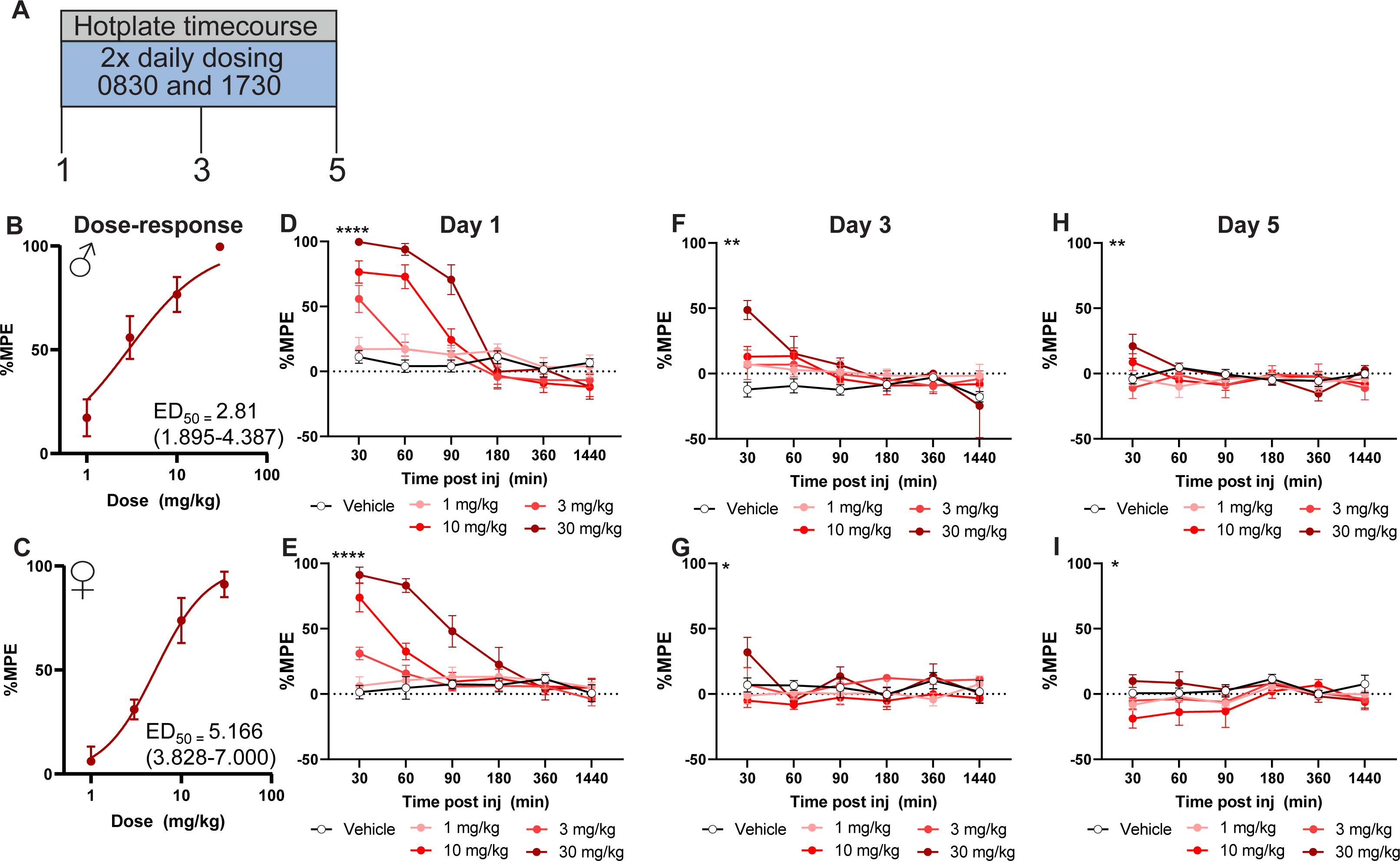
Oxycodone produces dose-dependent antinociception in the hotplate test. Oxycodone (1, 3, 10, 30 mg/kg s.c.) was administered twice daily over a period of 5 days (A). Oxycodone produced dose-dependent antinociception in males (B) and females (C) at the 30 min timepoint on day 1 of dosing. Tolerance developed by day 3 to most doses (Males: F, Females: G) and to all doses by day 5 (Males:H, Females:I). Data represented as mean ± SEM. N = 6-7 per group. ****p<0.0001, **p<0.01, *p<0.05 2×2 ANOVA interaction effect

Oxycodone produced dose-dependent antinociception that exhibited similar efficacy in both males (ED50 = 2.81 (1.90-4.39); Figure 1B) and females (ED50 = 5.166 (3.83-7.00); Figure 1C). Although the ED50 values are slightly higher in females, the 95% CI’s overlap indicating this difference is not significant. Similarly, Δ^9^-THC produced dose-dependent antinociception in both males (ED50 = 12.53 (8.24-19.28); Figure 2B) and females (ED50 = 19.20 (11.9-31.81); Figure 2C). Again, ED50 values in the females were larger however the CIs overlap indicating this difference is not significant.

**Figure 2.**
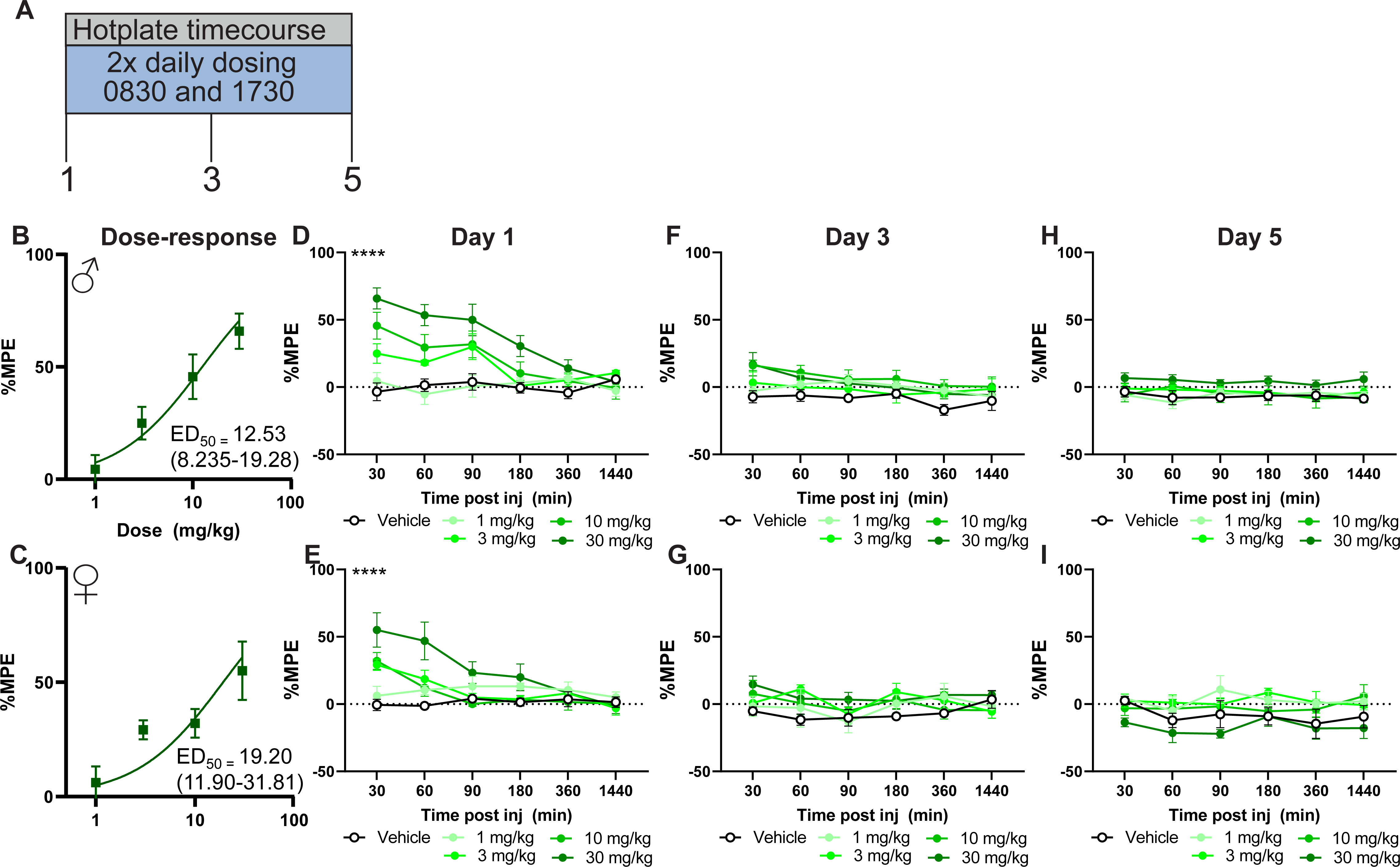
Δ^9^-THC produces dose-dependent antinociception in the hotplate test. Δ^9^-THC (1, 3, 10, 30 mg/kg s.c.) was administered twice daily over a period of 5 days (A). Δ^9^-THC produced dose-dependent antinociception in males (B) and females (C) at the 30 min timepoint on day 1 of dosing. Tolerance developed by day 3 (Males: F, Females: G) and remained by day 5 (Males:H, Females:I) in both sexes. Data represented as mean ± SEM. N = 6-7 per group. **** p<0.0001 2×2 ANOVA interaction effect

To determine the timecourse of oxycodone and Δ^9^-THC-induced antinociception, hotplate latencies were measured at 30, 60, 90, 180 and 1440 min post injection. At the highest dose tested (30 mg/kg s.c.) oxycodone-induced antinociceptive lasted until 90 min post injection in males (p<0.01 vs. vehicle two-way ANOVA followed by Tukey post-hoc; Figure 1D; Table 1), and 60 min post injection in females (p<0.01 vs. vehicle two-way ANOVA followed by Tukey post-hoc; Figure 1E; Table 1). Like oxycodone, Δ^9^-THC produced antinociception at 30 mg/kg s.c. dose for up to 90 mins post-injection in males (p<0.01 two-way ANOVA followed by Tukey’s post-hoc; Figure 2D; Table 2) but only 30 mins post-injection for females (p<0.01 two-way ANOVA followed by Tukey’s post-hoc; Figure 2E; Table 2)

By day 3 of dosing, antinociceptive tolerance had developed to all doses of oxycodone except the highest dose tested (30 mg/kg s.c.) in both males (Figure 1D; Table 1) and females (Figure 1E, Table 1). By day 5 all doses were ineffective relative to vehicle (Figure 1F,G). Δ^9^-THC - treated mice developed tolerance by day 3 in both males (Figure 2D) and females (Figure 2E) that remained on day 5 of dosing (Figure 2F,G, Table 2).

### A subthreshold dose of Δ^9^-THC enhances oxycodone-induced antinociception and delays the development of antinociceptive tolerance

Next, we sought to evaluate the effect of co-administration of Δ^9^-THC and oxycodone on hotplate antinociception and antinociceptive tolerance. To this end, we used the dose-response curves generated in Fig 1 to incorporate a dose of Δ^9^-THC which does not produce antinociception (3 mg/kg s.c.), combined with an effective, but not maximal antinociceptive dose of oxycodone (3 mg/kg s.c.). Animals received either oxycodone (3 mg/kg s.c.), Δ^9^-THC (3 mg/kg s.c.) a combination of oxycodone (3 mg/kg s.c.) and Δ^9^-THC (3 mg/kg s.c.) or vehicle twice daily for a period of five days (see Figure 3A for timeline). Combination groups exhibited enhanced antinociception relative to all groups on day 1 and at the 30 min timepoint (Males: F15, 80= 6.968, p<0.01, Females: F15, 80= 5.596, p<0.01 two-way ANOVA interaction effect; Figure 3B,C; Table 3) and on day 3 at the 30 min timepoint (Males: F15, 80= 4.194, p<0.01, Females: F15, 80= 3.286, p<0.01 two-way ANOVA interaction effect; Figure 3D,E;Table 3). However, by day 5 there were no differences between groups (Figure 3F,G). These data suggest that a subthreshold dose of Δ^9^-THC can augment oxycodone-induced antinociception.

**Figure 3.**
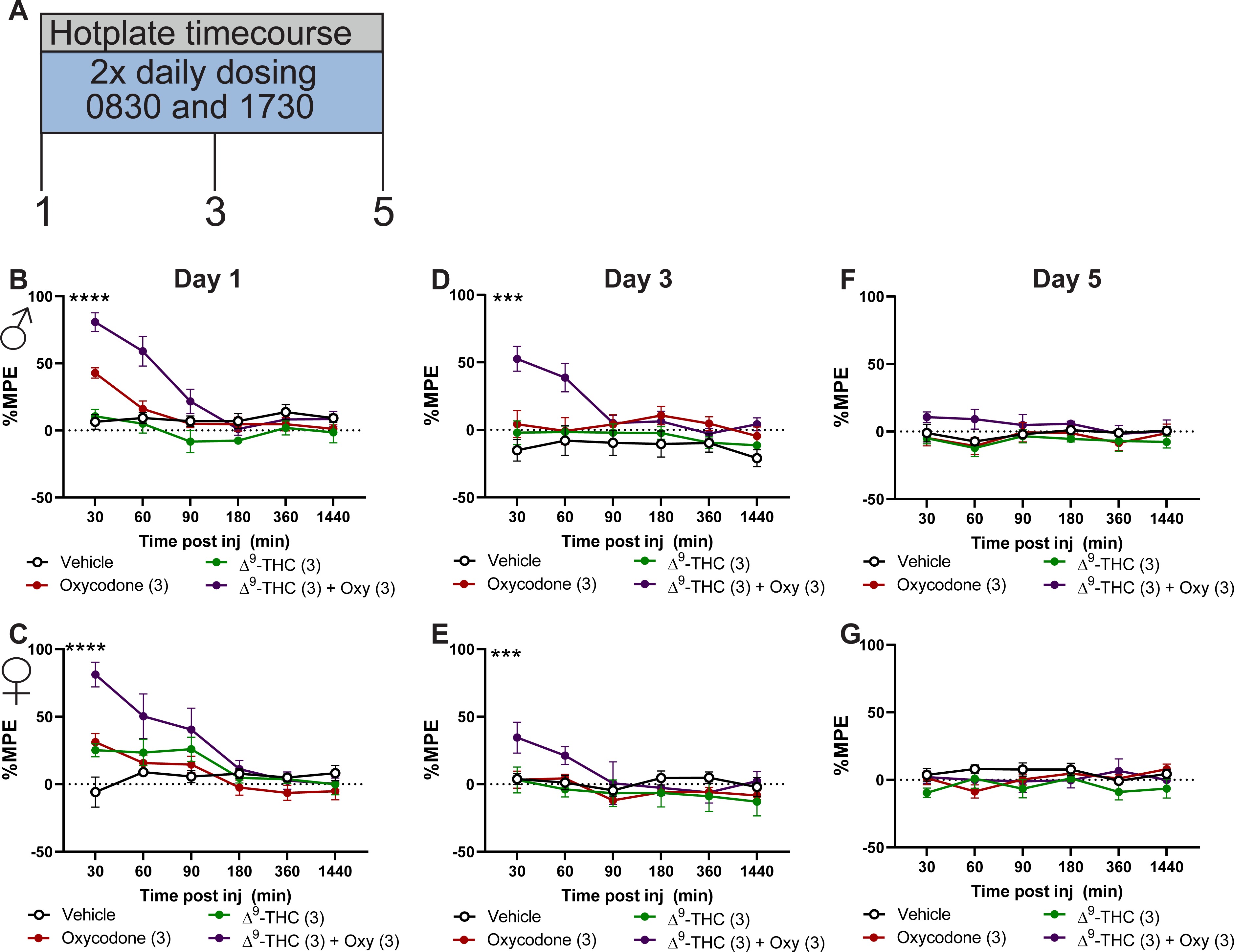
Combination dosing of Δ⁹-THC with oxycodone results in enhanced antinociception and delayed antinociceptive tolerance. Schematic of experimental timecourse (A). Oxycodone (3 mg/kg s.c.) and Δ^9^-THC (3 mg/kg s.c.) produced greater and longer lasting antinociception when co-administered relative to either administered alone on day 1 of dosing in both males (B) and females (C). On day 3 of dosing, the combination remained efficacious, but tolerance had developed to the other treatment groups in both males (D) and females (E). By day 5 complete tolerance had developed in all groups in both males (F) and females (G). Data represented as mean ± SEM. N = 5 per group. **** p<0.0001, *** p<0.001 two-way ANOVA interaction effect.

### Repeated dosing with oxycodone produces physical dependence. Combination treatment of Δ^9^-THC and oxycodone reduced physical oxycodone-induced physical dependence in male but not female mice

We next sought to evaluate if our repeated dosing regimen would produce µ-opioid receptor dependence. Repeated administration of µ-opioid agonists produces physical dependence that can be precipitated with administration of naloxone. Mice were administered naloxone (3 mg/kg s.c.) on day 6 of repeated dosing and evaluated for jumping behavior over 30 mins. Male (F4,24= 5.039, p<0.01 one-way ANOVA; Figure 4B;Table 4) and female (F4,24= 4.692, p<0.01 one-way ANOVA; Figure 4C; Table 4) mice treated with oxycodone exhibited jumping behavior following naloxone treatment, indicative of µ-opioid mediated dependence. Combination treatment with Δ^9^-THC reduced naloxone-precipitated jumping behavior in males (F3,16= 7.853, p<0.01 one-way ANOVA; Figure 4D; Table 4) but not females (F3,16= 5.300, p<0.01 one-way ANOVA; Figure 4E; Table 4). These data indicate that oxycodone administration produces µ-opioid receptor dependence, and Δ^9^-THC administration does not augment this effect.

**Figure 4.**
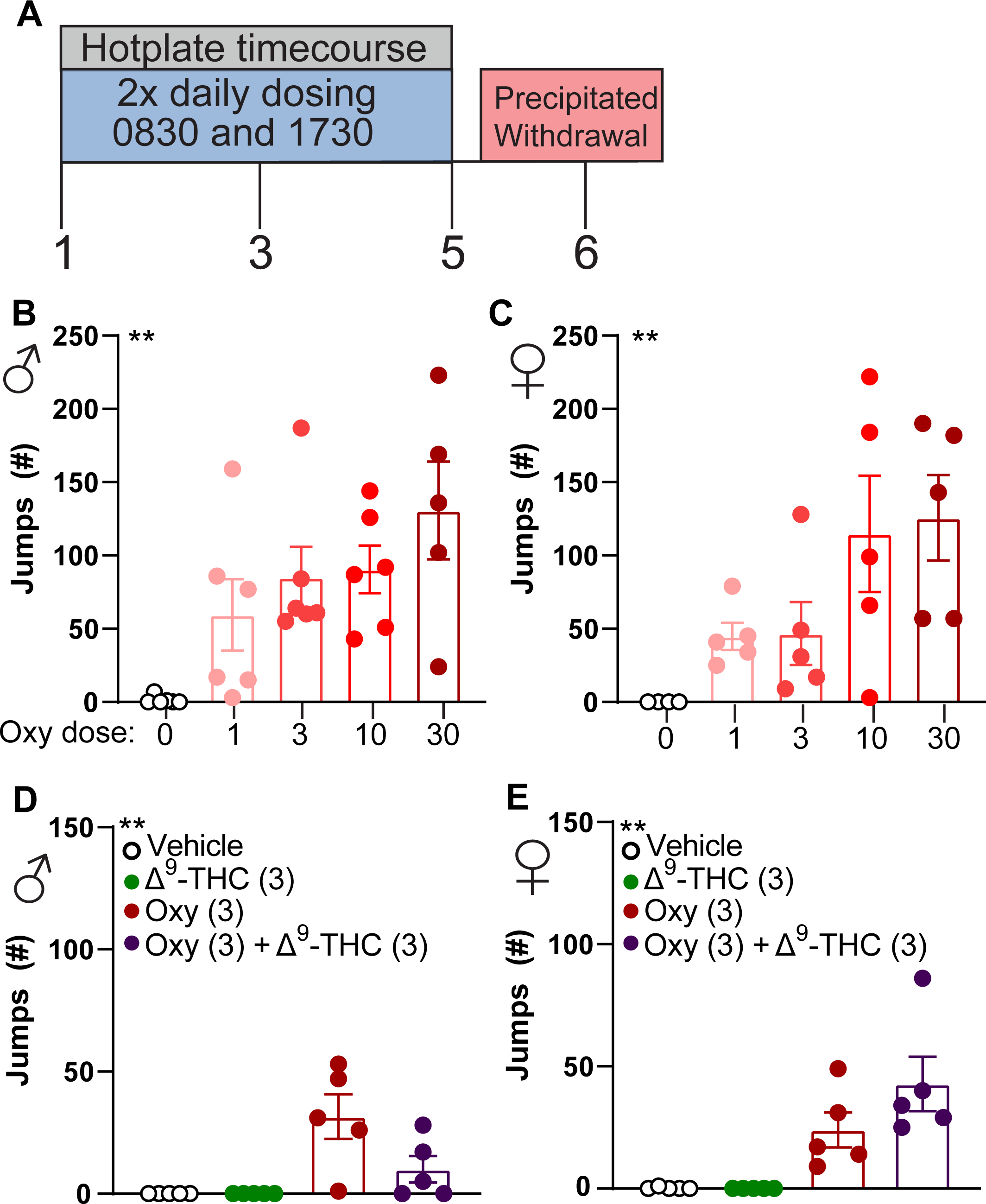
Repeated dosing with oxycodone induces µ-opioid receptor dependence, co-administration of oxycodone with Δ^9^-THC reduces µ-opioid receptor dependence in males. Oxycodone produced dependence to a similar degree in both males (B) and females (C) following repeated dosing. Males treated with Δ^9^-THC and oxycodone in combination had a reduced number of jumps (D), whereas the females did not differ from oxycodone alone (E). Data are represented as mean ± SEM. N = 5-7 per group. **p<0.01 one-way ANOVA main effect.

### Treatment with oxycodone or a combination of oxycodone and Δ^9^-THC results in alterations of circadian activity

Repeated opioid intake can cause alterations in circadian activity(Stinus et al., 1998; Bedard et al., 2023). Insomnia is also a hallmark sign of withdrawal from opioids(Budney et al., 2004) and cannabis(Budney et al., 2004; Babson et al., 2013a; Bonn-Miller et al., 2019). We sought to evaluate if our dosing regimen with oxycodone, Δ^9^-THC or a combination would produce changes in general circadian activity during dosing, withdrawal, and recovery periods (See Figure 5A for timeline). Mice treated with Δ^9^-THC (3 mg/kg s.c.) did not exhibit alterations in circadian activity relative to vehicle-treated controls in either males (Figure 5B,H; Table 5) or females (Figure 5C,I; Table 5). Mice treated with oxycodone (3 mg/kg s.c.) were more active during the dosing period relative to vehicle treated controls in both sexes (Male: two-way ANOVA interaction effect; Figures D,H,E,I). Combination treatment of Δ^9^-THC (3 mg/kg s.c.) and oxycodone (3 mg/kg s.c.) resulted in increased activity during the light period in both sexes (p<0.01 two-way ANOVA followed by Tukey post-hoc). See Supplemental figure 1 for acute activity increase, Supplemental Figure 2 for increase throughout the dosing phase. Total activity in the light period was normalized during the 2-day recovery period following withdrawal. These data indicate that oxycodone can produce detectable alterations in circadian activity, and co-treatment with Δ^9^-THC does not affect oxycodone-induced alterations in circadian activity. See supplemental figure 2 for circadian activity traces throughout all phases of the experiment.

**Figure 5.**
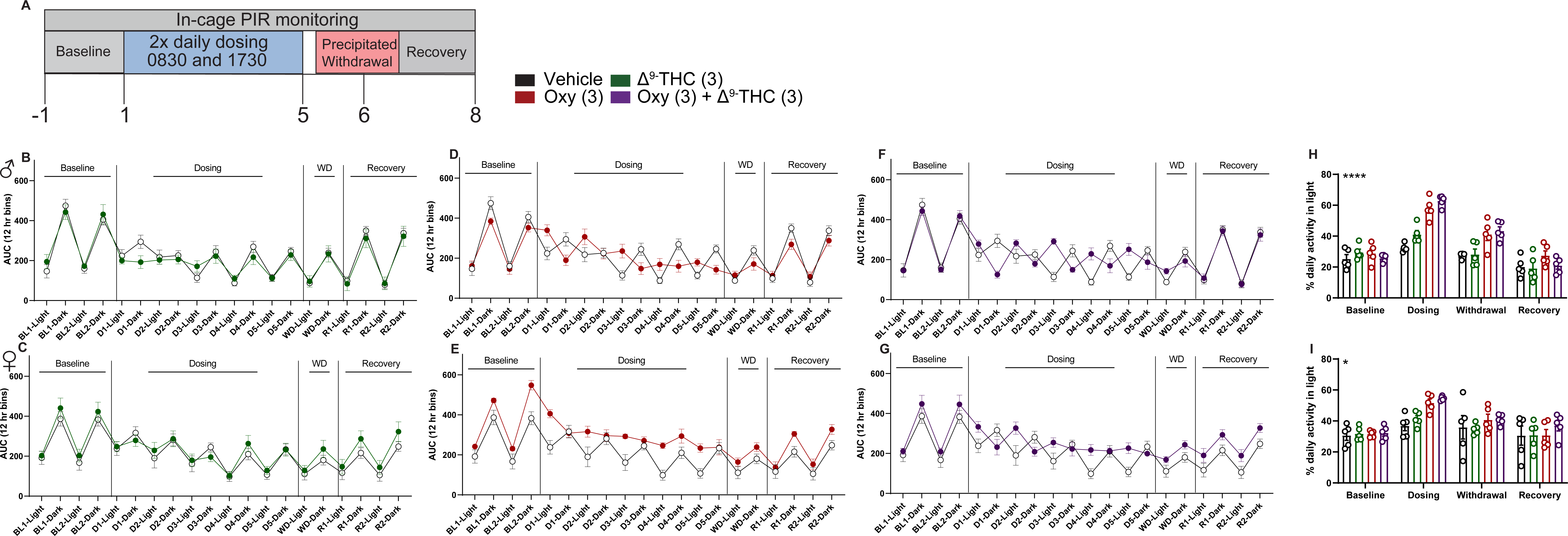
Dosing with oxycodone or a combination of Δ^9^-THC with oxycodone results in disturbances of circadian activity. Δ^9^-THC had a minimal effect on general circadian activity pattern relative to vehicle (B,C). Mice treated with oxycodone (D,E) or a combination of oxycodone+Δ^9^-THC (F,G) spent a greater percentage of time active in the light relative to vehicle during the dosing period which normalized after withdrawal in both males (H) and females (I). Values for percent of time in light were averaged across each experimental phase and compared in panels D and H. Data represented as mean ± SEM. N = 5 per group **** p<0.0001, *p<0.05 2×2 ANOVA interaction effect.

### Δ^9^-THC facilitates oxycodone-induced CPP and does not alter oxycodone-induced locomotor sensitization

Δ^9^-THC has been demonstrated to enhance the subjective effects of oxycodone in humans(Cooper et al., 2018). However, Δ^9^-THC can reduce oxycodone self-administration in rats(Nguyen et al., 2019) and morphine-induced CPP is reduced in CB1 knockout mice(Martin et al., 2000; Iyer et al., 2022). Here we incorporated CPP to evaluate if Δ^9^-THC would enhance or dampen oxycodone-induced CPP at doses therapeutically relevant for antinociception.

Mice administered vehicle for pairing conditions did not exhibit changes in chamber preference (Supplemental Figure 3A,B). Δ^9^-THC did not produce CPP in either sex (Supplemental Figure C,D). Mice administered oxycodone at 1 mg/kg s.c. did not exhibit CPP for the drug-paired chamber (Figure 6A,B). In contrast, mice treated with a 3 mg/kg s.c. dose of oxycodone developed a robust CPP to the drug-paired chamber (Male: p<0.01, Female: p<0.01 vs. vehicle paired-chamber two-way ANOVA followed by Bonferroni post-hoc; Figure 6C,D; Table 6).

**Figure 6.**
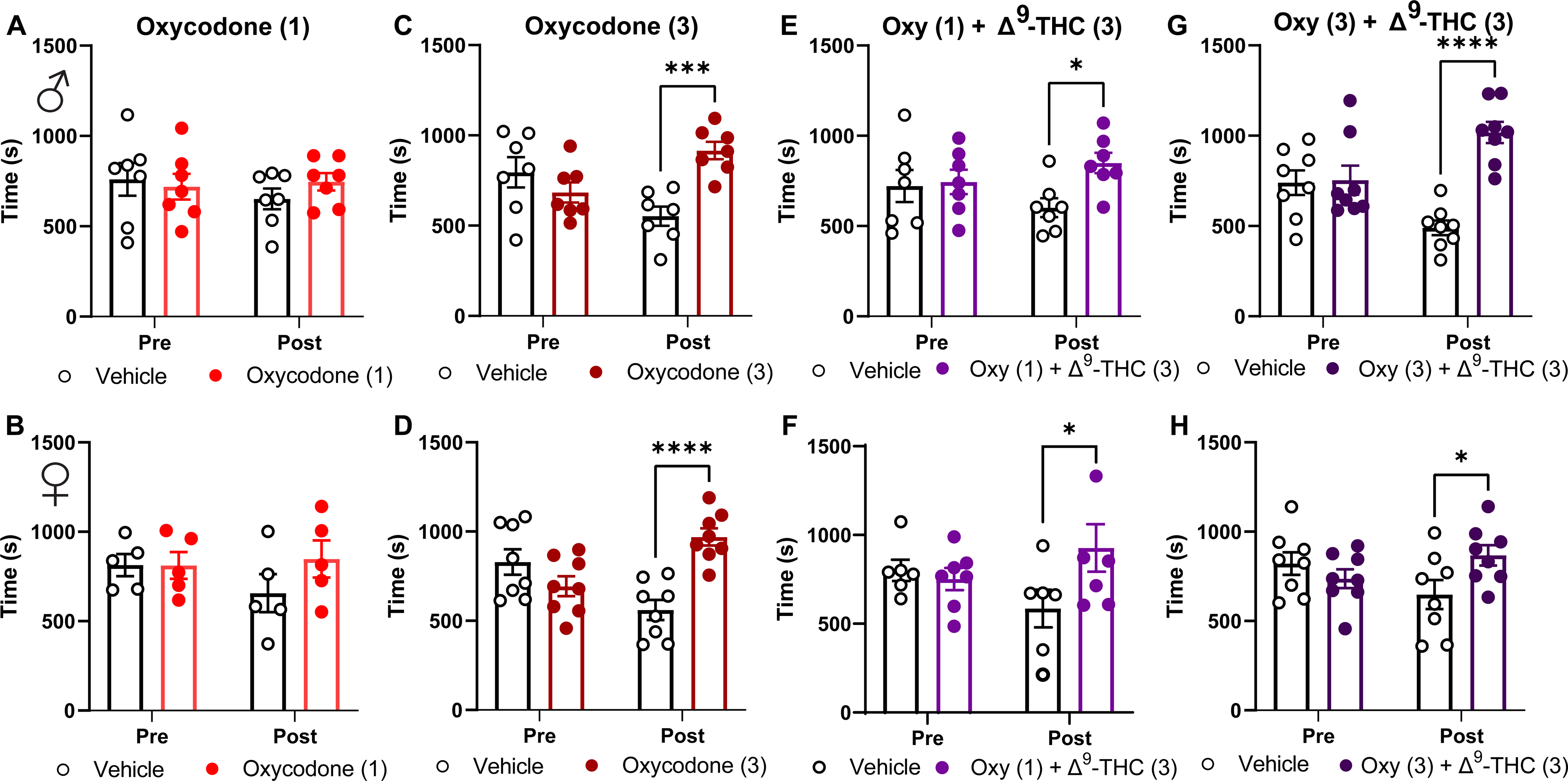
Oxycodone and Δ^9^-THC produce conditioned place preference when administered in combination. Administration of oxycodone (1 mg/kg s.c.) did not produce CPP in males (A) or females (B). By contrast, a 3 mg/kg dose of oxycodone produced CPP in both males (C) and females (D). Doses of Δ^9^-THC (3 mg/kg s.c.) and oxycodone (1 mg/kg s.c.) that did not produce preference on their own produced CPP when co-administered in both males (E) and females (F). Δ^9^-THC (3 mg/kg s.c.) did not alter oxycodone (3 mg/kg s.c.)-induced CPP in either males (G) or females (H). Data represented as mean ± SEM. N = 5-8 per group ****p<0.0001, *** p<0.001, *p<0.05 2×2 ANOVA followed by Sidak post-hoc.

We next evaluated if doses of oxycodone (1 mg/kg s.c.) and Δ^9^-THC (3 mg/kg s.c.), which were subthreshold for producing preference alone, would produce preference when co-administered. Mice administered this combination produced CPP in both sexes (Male: p<0.04, Female: p<0.03 vs. vehicle-paired chamber two-way ANOVA followed by Sidak post-hoc; Figure 6C,D; Table 6). Similarly, combination of a preference producing oxycodone dose (3 mg/kg s.c.) and Δ^9^-THC (3 mg/kg s.c.) produced CPP (Male: p<0.01, Female: p<0.05 two-way ANOVA followed by Sidak post-hoc; Figure 6 E,F; Table 6). These data indicate that Δ^9^-THC does not reduce oxycodone-induced CPP and subthreshold doses of oxycodone and Δ^9^-THC become preference producing when combined.

µ-opioid agonists produce an initial hyperlocomotor response that sensitizes following repeated administration(Severino et al., 2020), which parallels the development of drug-seeking behavior(Robinson and Berridge, 1993; Brings et al., 2022). We tracked locomotion during the CPP pairing days to evaluate this effect across our different groups. Oxycodone produced increased locomotion at 3 mg/kg in both sexes across all pairing days (Male: F5,39= 43.00, p<0.01, Female: F5,29= 12.29 p<0.01 two-way ANOVA treatment effect; Figure 7A,B; Table 7). The combination group of oxycodone (3 mg/kg s.c.) with Δ^9^-THC (3 mg/kg s.c.) also produced enhanced locomotion relative to all other groups aside from the oxycodone (3 mg/kg s.c.) group in both sexes. Although the low dose oxycodone (1 mg/kg s.c.) and Δ^9^-THC (3 mg/kg s.c.) combination group produced preference there was no increase in locomotion or indication of locomotor sensitization in either sex. These data indicate that while Δ^9^-THC can facilitate oxycodone-induced CPP, it does not alter oxycodone-induced locomotor sensitization.

**Figure 7:**
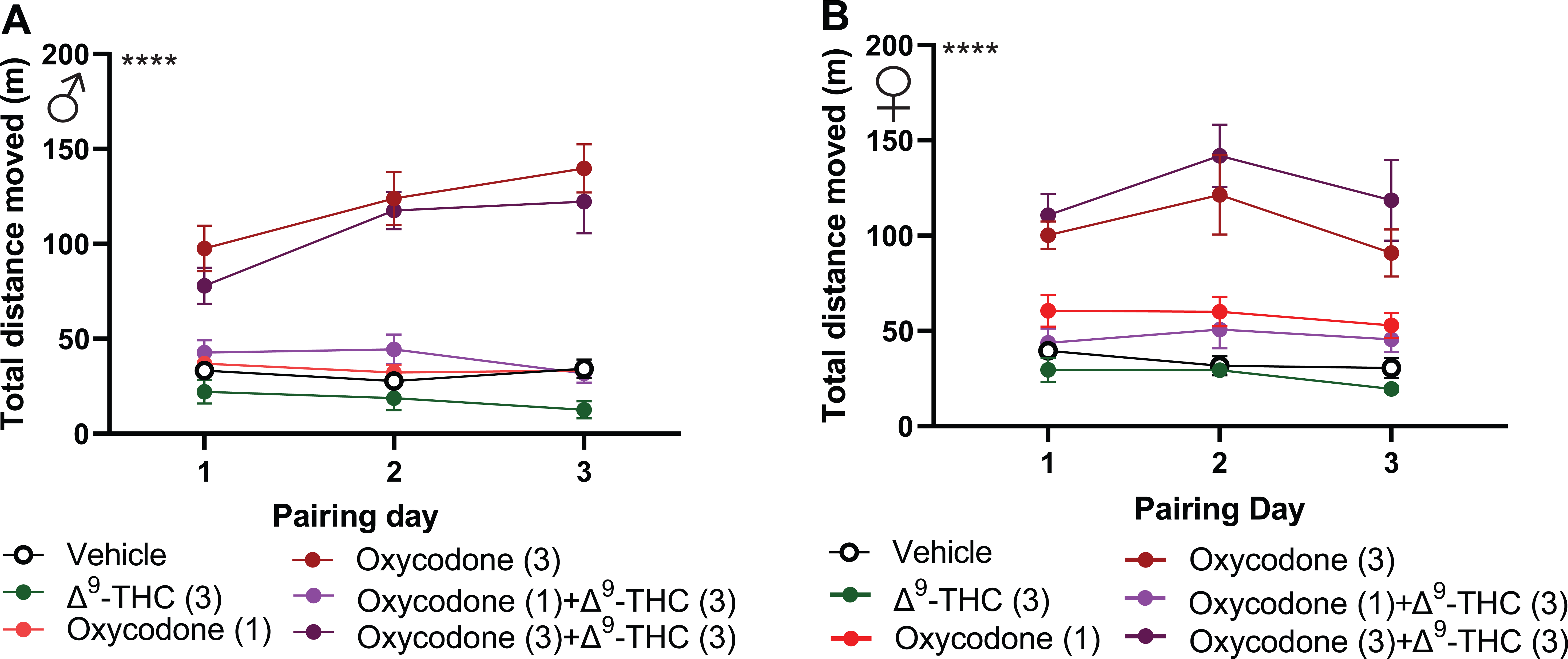
Δ^9^-THC does not alter oxycodone-induced locomotor sensitization. Distance traveled was measured during each pairing day from during CPP conditioning. Oxycodone increased distance travelled, while Δ^9^-THC had no effect in both males (A) and females (B) Δ^9^-THC also did not alter distance traveled when co-administered with oxycodone. Data represented as mean ± SEM. N = 5-8 per group. **** p<0.0001 2×2 ANOVA interaction effect.

## Discussion

In the present study we demonstrate that combination treatment of Δ^9^-THC and oxycodone results in greater antinociceptive efficacy and delayed antinociceptive tolerance relative to either administered alone. Using doses which produce antinociception, we demonstrate that Δ^9^-THC does not alter oxycodone-induced µ-opioid receptor dependence or alterations in circadian activity. We also demonstrate that co-treatment of oxycodone and Δ^9^-THC produces CPP at doses in which either in isolation do not produce CPP. These results have therapeutic implications for combination treatment of oxycodone and Δ^9^-THC.

Both oxycodone and Δ^9^-THC produced dose-dependent antinociception in the hotplate. Prior studies have reported that females are more sensitive to the antinociceptive effects of Δ^9^-THC(Craft et al., 2013; Wakley et al., 2014), and have demonstrated differential rates of tolerance development in the context of analgesia(Henderson-Redmond et al., 2022; Piscura et al., 2023). In our study, Δ^9^-THC produced a higher ED50 in females relative to males, but this was not statistically significant. This is consistent with a prior report indicating a higher ED50 for Δ^9^-THC in the tail-flick test in mice(Henderson-Redmond et al., 2022). Yet, these data are in contrast to other reports in rats which suggest females are more sensitive(Tseng and Craft, 2001; Wakley and Craft, 2011a; Craft et al., 2012; Wakley et al., 2014)(for review see:(Cooper and Craft, 2018)) and reflect a species difference. We also did not observe a sex difference in oxycodone-induced analgesia. This is in parallel to prior reports indicating similar antinociceptive efficacy of oxycodone between sexes(Collins et al., 2016). Tolerance developed to repeated administration in mice treated with either compound and was similar between sexes. When Δ^9^-THC was co-administered at a sub-ED50 dose (3 mg/kg s.c.) with oxycodone at a near-ED50 dose (3 mg/kg s.c.), the effect of the combination was greater than either administered alone. Interestingly, we observed a delay in the development of tolerance in animals administered the combination relative to either drug alone in both sexes. This is similar to what others have reported using Δ^9^-THC in combination with oxycodone and other opioids such as morphine on the tail-flick test in rodents(Welch and Stevens, 1992; Cichewicz et al., 1999; Yesilyurt et al., 2003; Williams et al., 2006; Wakley and Craft, 2011b)(for review see (Nielsen et al., 2022)). Such enhancements of antinociceptive efficacy are also observed in nonhuman primates(Beardsley et al., 2004; Maguire and France, 2014; Gerak and France, 2016; Carey et al., 2023) and in some cases, humans(Cooper et al., 2018). Excitingly there are two clinical studies currently ongoing investigating interactions of Δ^9^-THC and oxycodone in an analgesic context. One in the context of fibromyalgia pain(van Dam et al., 2023) and another in a naïve context using the cold pressor test (Clinicaltrials.gov identifier: NCT03679949). It will be interesting to see if an enhanced analgesic effect is observed in the combination groups of these trials.

Prior reports demonstrate that direct CB1 agonists, allosteric modulators, and endocannabinoid degradation inhibitors can reduce somatic and autonomic withdrawal in opioid-dependent animals(Scavone et al., 2013). Further, co-treatment of Δ^9^-THC with fentanyl does not alter opioid withdrawal signs in nonhuman primates(Gerak and France, 2016). These lines of evidence suggest Δ^9^-THC likely wouldn’t facilitate and might reduce withdrawal-associated behaviors from oxycodone. Like prior reports(Enga et al., 2016; Carper et al., 2021), naloxone induced jumping behavior in mice treated repeatedly with oxycodone. Notably, jumping behavior was present even at the lowest dose tested(1 mg/kg s.c.), indicating does that do not exhibit antinociceptive efficacy are still sufficient to produce dependence. In males, co-administration with Δ^9^-THC and oxycodone reduced jumping, however this effect was not observed in females. These results suggest that Δ^9^-THC did not exacerbate µ-opioid receptor physical dependence at doses relevant for facilitating antinociception and may even reduce opioid withdrawal signs..

Alterations in sleep and circadian rhythm can occur following repeated intake of and withdrawal from opioids and cannabis(Beswick et al., 2003; Levin et al., 2010; Vandrey et al., 2011; Babson et al., 2013b; Gorelick et al., 2013; Eacret et al., 2020; Tripathi et al., 2020; Tamura et al., 2021; Bergum et al., 2022; Gamble et al., 2022; Xue et al., 2022). To our knowledge, our study is the first to evaluate combination treatment of oxycodone and Δ^9^-THC in such metrics. We report that twice daily administration of oxycodone(3 mg/kg s.c.) resulted in a profound impact on circadian activity which was normalized following withdrawal in both sexes. Other studies have demonstrated a longer-lasting impact on circadian activity and sleep behavior following withdrawal from morphine(Marchant and Mistlberger, 1995; Stinus et al., 1998) or fentanyl(Gamble et al., 2022). These studies did not precipitate withdrawal, and incorporated opioid ligands and dosing regimens which may account for the lack of a continued withdrawal effect in the present study. Regardless, there are still apparent alterations in activity that are induced by oxycodone dosing, which parallels our prior report on voluntary oral consumption of oxycodone in mice, where mice consuming oxycodone exhibited increased activity during the light phase(Slivicki et al., 2023). Δ^9^-THC, in contrast, did not appear to affect circadian activity during any phase of the observation period. This differs from recent studies which demonstrate alterations in sleep patterns following administration of Δ^9^-THC(Kesner et al., 2022) or administration of a synthetic cannabinoid agonist, AM2389(Missig et al., 2022). The cause of such differences may be the endpoints used (EEG vs. cage activity), larger dose of Δ^9^-THC (10 mg/kg) or different CB1 agonist (AM2389). Δ^9^-THC did not appear to alter oxycodone’s impact on circadian activity during any point of the study. A limitation of this study is that drug administration occurred during the animal’s light cycle when they generally have less activity, and using a dose of oxycodone that produces hyperlocomotion.

Prior reports have demonstrated interactions between cannabinoids and opioids on measures of reward in rodents(Iyer et al., 2022; Mohammadkhani and Borgland, 2022), nonhuman primates(Beardsley et al., 2004; Maguire et al., 2013; Gerak et al., 2019) and humans(Cooper et al., 2018). It is therefore important to consider reward-related measures of Δ^9^-THC and oxycodone in a dose range that is therapeutically relevant. In our study, oxycodone produced dose-dependent CPP in both sexes at the 3 mg/kg dose. This is consistent with prior literature demonstrating CPP in mice using this dose(Niikura et al., 2013; Harris et al., 2022). By contrast Δ^9^-THC did not produce conditioned place preference at the 3 mg/kg s.c. dose. When non-preference producing doses of Δ^9^-THC (3 mg/kg) and oxycodone (1 mg/kg) were administered, it resulted in the development of CPP in both sexes. These findings parallel recent literature reporting that vaporized or injected Δ^9^-THC reduces oxycodone(Nguyen et al., 2019; Nguyen et al., 2021; Nguyen et al., 2023) and heroin(Gutierrez et al., 2022) self-administration in rats. The authors in(Nguyen et al., 2021; Nguyen et al., 2023) suggest that Δ^9^-THC may be increasing the rewarding efficacy of a “unit dose” of oxycodone when self-administered. Our findings in the CPP paradigm are consistent with this hypothesis. Furthermore, smoked cannabis produced an increase in oxycodone-induced drug liking and subjective effects in humans(Lofwall et al., 2016; Cooper et al., 2018). This interaction has the potential to be detrimental in a clinical setting. Co-administering Δ^9^-THC with oxycodone could result in a greater association of reward or liking with oxycodone, potentially resulting in an increased likelihood of drug seeking behavior. On the other hand, Δ^9^-THC could be used as an adjuvant therapy for dose tapering in the case of oxycodone dependence or in combination with other opioid-based therapies such as methadone(Strang et al., 2020). Regardless, there are clear interactions between Δ^9^-THC and oxycodone in reward and drug “liking” across species that warrant consideration with respect to therapeutic implementation of combination treatments.

Locomotor sensitization following repeated opioid exposure has been implicated as a proxy for drug seeking behavior(Robinson and Berridge, 1993; Valjent et al., 2010). CB1 agonists can induce catalepsy which may obscure interpretation of locomotor-based assays(Compton et al., 1993; Metna-Laurent et al., 2017; Slivicki et al., 2022). Δ^9^-THC did not reduce locomotor activity at the dose tested here in either sex, allowing us to study the effect of Δ^9^-THC on endpoints of oxycodone-induced locomotor activity. A prior report in rats found that pretreatment with Δ^9^-THC (41 days) increased heroin-induced locomotor sensitization(Singh et al., 2005). CB1 knockout mice do not exhibit locomotor sensitization when administered morphine(Martin et al., 2000). In contrast to these results our data indicate that co-administration of Δ^9^-THC with oxycodone does not alter oxycodone-induced hyperlocomotion or locomotor sensitization. This may reflect difference in the ligands and/or dosing conditions used.

The findings of the present study complement and extend the existing literature on cannabinoid and opioid interactions. Taken together, our results support the notion that the combination of Δ^9^-THC and oxycodone may result in a facilitation of desired therapeutic effects (e.g. antinociception, delayed tolerance) without augmenting certain undesired effects of µ-opioid receptor activation (e.g. dependence and circadian rhythm alteration). However, our data also suggests the abuse liability of such combinations warrants consideration. Future studies evaluating such metrics in the context of pain will be of great importance to fully comprehend the clinical utility of cannabinoid and opioid combination therapies.

## Supporting information

Statistics Tables

## Acknowledgments

Grant support from F32DA051160 (to RAS) R34NS126036 (RWG), R01DA049924 (MCC), R01DA058755 (MCC), Hetzler Foundation for Addiction Research and Prevention (MCC), Diabetes Research Center Pilot Project (AVK) Hope Center Pilot Project (to AVK) Whitehall Foundation Grant (2017-12-54 to MCC) Rita Allen Scholar Award in Pain (to MCC). This work was also supported by the Dr. Seymour and Rose T. Brown Professorship in Anesthesiology (RWG).

---

**Supplemental Figure 1.**
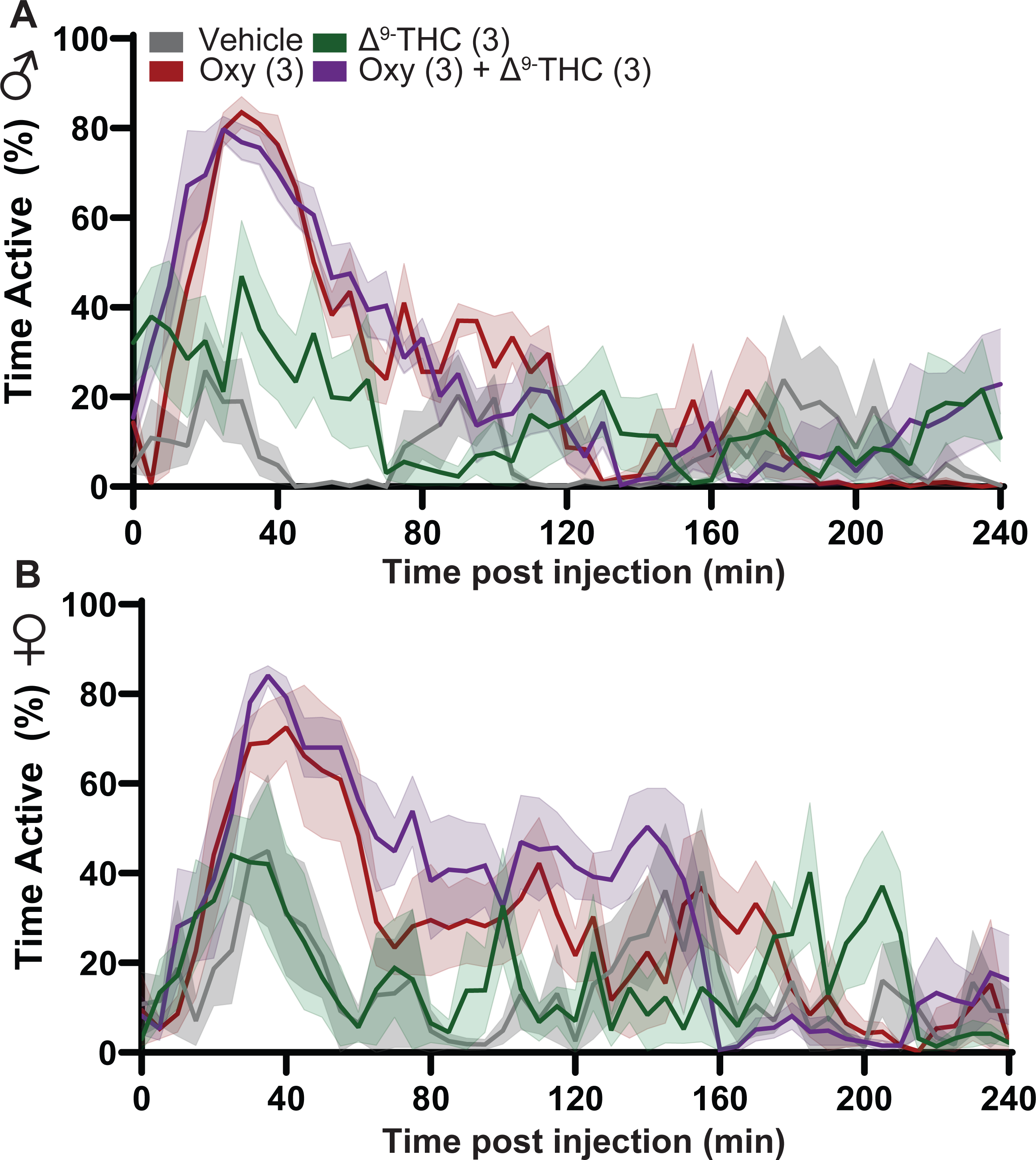
Homecage activity traces following a drug administration. Activity data from Pallidus devices were plotted as 5 min bins. Vehicle, Oxycodone (3 mg/kg s.c.), Δ^9^-THC (3 mg/kg s.c.), or a combination of oxycodone and Δ^9^-THC was administered at the 0 min mark in both males (A) and females (B). Oxycodone and combination treatment of oxycodone and Δ^9^-THC produced transient increases in activity in both males and females. Data represented as mean ± SEM. N = 5-6 per group

**Supplemental Figure 2.**
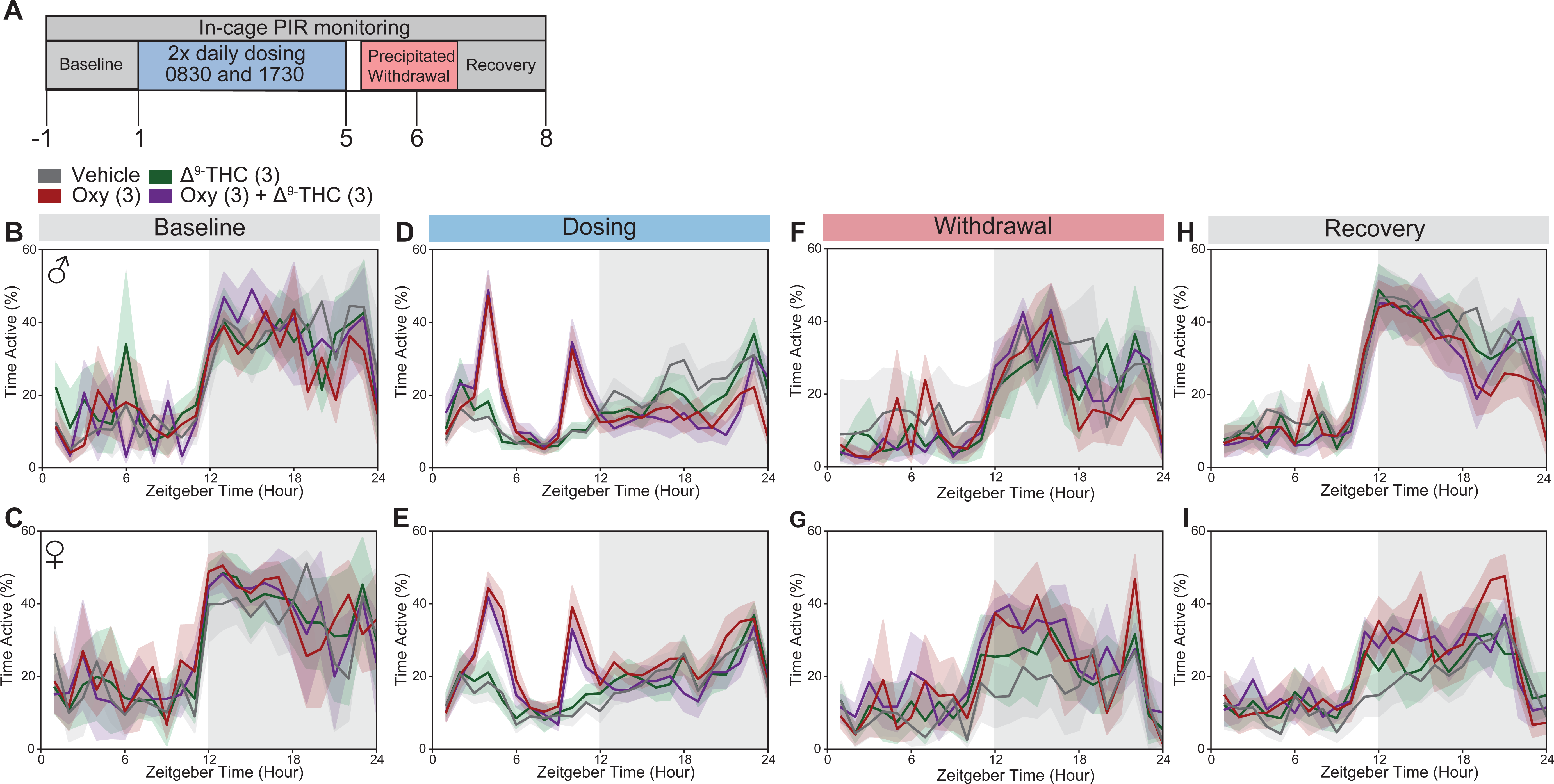
Oxycodone and combination of Δ^9^-THC with oxycodone produce alterations in circadian activity during the dosing phase. Schematic of experimental timecourse (A). Activity data from Pallidus devices were plotted as 1 hr bins and averaged for each phase of the experiment. Activity was similar across all groups during the baseline phase of the experiment in both males (B) and females (C). During the dosing phase, treatment with oxycodone and a combination of oxycodone and Δ^9^-THC produced a spike in activity during the light phase in both males (D) and females (E) that corresponded to the time when the treatment was administered. There were slight differences between groups during the withdrawal phase (F,G) and groups appeared mostly similar during the recovery phase (H,I). Data represented as mean ± SEM. N = 5 per group

**Supplemental Figure 3.**
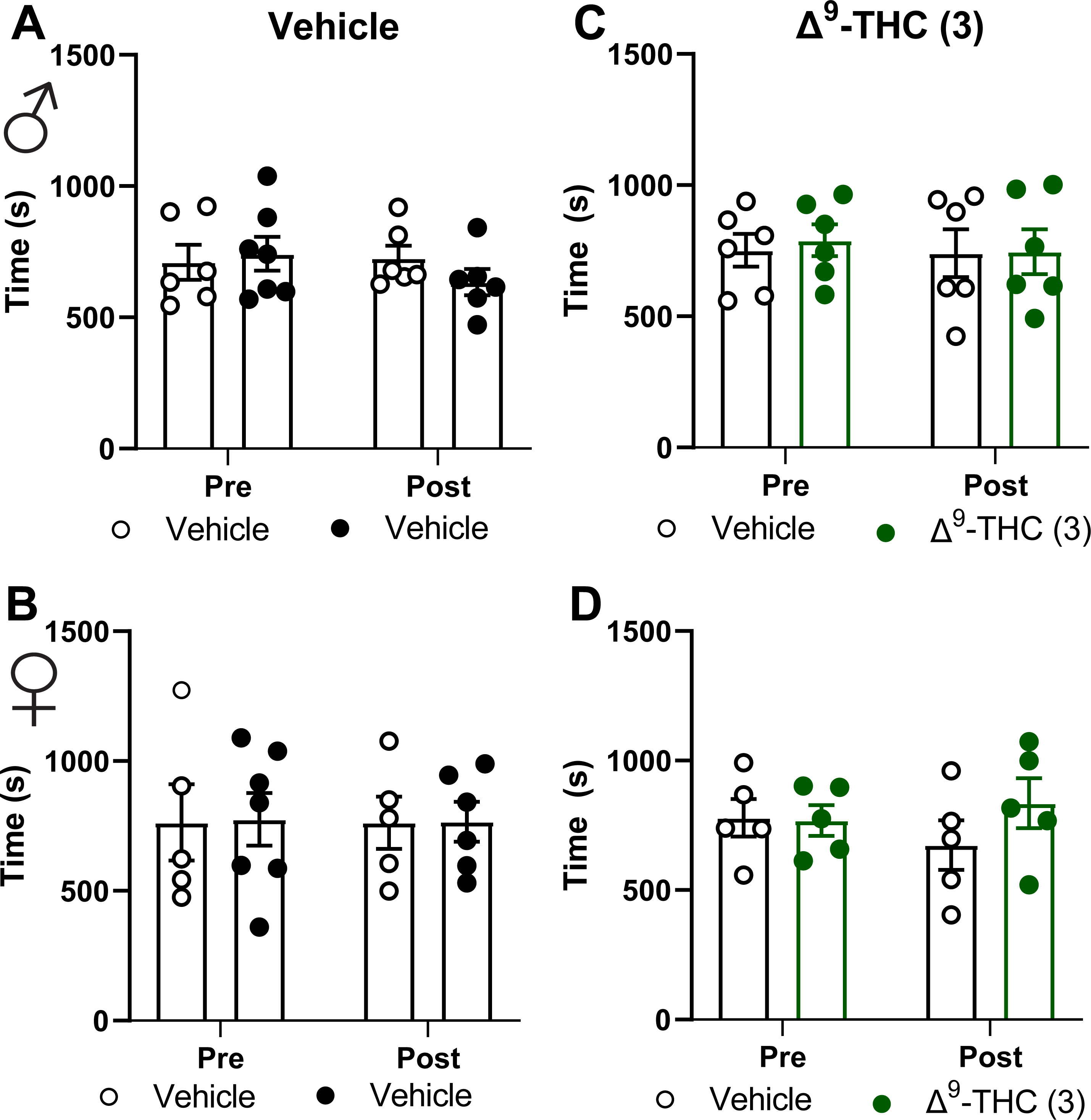
Vehicle and Δ^9^-THC do not produce alterations in chamber preference. Administration of vehicle only or Δ^9^-THC (3 mg/kg s.c.) did not produce alterations in chamber preference in either males (A,C) or females (B,E). Data represented as mean ± SEM. N = 5-6 per group

