## Supplementary material for "Impact of Δ^9^-Tetrahydrocannabinol and oxycodone co-administration on measures of antinociception, dependence, circadian activity, and reward in mice": Statistics Tables

**Table 1. Statistics from Figure 1**

| **Figure 1D – Male Day 1** | | | |
| --- | --- | --- | --- |
| **Type of test** | **Comparison** | **P-value summary** | **F, P-value** |
| Two-way ANOVA | Time | **** | F _(3.548, 88.70)_ = 56.29, P<0.0001 |
|  | Treatment | **** | F _(4, 25)_ = 15.83, P<0.0001 |
|  | Interaction | **** | F _(20,125)_ = 10.41, P<0.0001 |
| Tukey’s post-hoc |  |  |  |
| 30 | Vehicle vs. 1 mg/kg | ns | 0.9732 |
|  | Vehicle vs. 3 mg/kg | * | 0.0332 |
|  | Vehicle vs. 10 mg/kg | *** | 0.001 |
|  | Vehicle vs. 30 mg/kg | **** | <0.0001 |
|  | 1 mg/kg vs. 3 mg/kg | ns | 0.105 |
|  | 1 mg/kg vs. 10 mg/kg | ** | 0.0049 |
|  | 1 mg/kg vs. 30 mg/kg | ** | 0.0013 |
|  | 3 mg/kg vs. 10 mg/kg | ns | 0.5593 |
|  | 3 mg/kg vs. 30 mg/kg | * | 0.0415 |
| 60 | 10 mg/kg vs. 30 mg/kg | ns | 0.1816 |
|  | Vehicle vs. 1 mg/kg | ns | 0.8172 |
|  | Vehicle vs. 3 mg/kg | ns | 0.3981 |
|  | Vehicle vs. 10 mg/kg | ** | 0.0013 |
|  | Vehicle vs. 30 mg/kg | **** | <0.0001 |
|  | 1 mg/kg vs. 3 mg/kg | ns | >0.9999 |
|  | 1 mg/kg vs. 10 mg/kg | * | 0.0245 |
|  | 1 mg/kg vs. 30 mg/kg | ** | 0.0036 |
|  | 3 mg/kg vs. 10 mg/kg | ** | 0.005 |
|  | 3 mg/kg vs. 30 mg/kg | **** | <0.0001 |
|  | 10 mg/kg vs. 30 mg/kg | ns | 0.3323 |
| 90 | Vehicle vs. 1 mg/kg | ns | 0.7073 |
|  | Vehicle vs. 3 mg/kg | ns | 0.8368 |
|  | Vehicle vs. 10 mg/kg | ns | 0.3137 |
|  | Vehicle vs. 30 mg/kg | * | 0.0131 |
|  | 1 mg/kg vs. 3 mg/kg | ns | >0.9999 |
|  | 1 mg/kg vs. 10 mg/kg | ns | 0.7778 |
|  | 1 mg/kg vs. 30 mg/kg | * | 0.0238 |
|  | 3 mg/kg vs. 10 mg/kg | ns | 0.8368 |
|  | 3 mg/kg vs. 30 mg/kg | * | 0.022 |
|  | 10 mg/kg vs. 30 mg/kg | ns | 0.0676 |
| **Figure 1E – Female day 1** | | | |
| **Type of test** | **Comparison** | **P-value summary** | **F, P-value** |
| Two-way ANOVA | Time | **** | F _(3.075, 61.51)_ = 56.29, P<0.0001 |
|  | Treatment | **** | F _(4, 25)_ = 13.42, P<0.0001 |
|  | Interaction | **** | F _(20,100)_ = 10.18, P<0.0001 |
| Tukey’s post-hoc |  |  |  |
| 30 | Vehicle vs. 1 mg/kg | ns | 0.9807 |
|  | Vehicle vs. 3 mg/kg | * | 0.0179 |
|  | Vehicle vs. 10 mg/kg | ** | 0.0063 |
|  | Vehicle vs. 30 mg/kg | **** | <0.0001 |
|  | 1 mg/kg vs. 3 mg/kg | ns | 0.1141 |
|  | 1 mg/kg vs. 10 mg/kg | ** | 0.0078 |
|  | 1 mg/kg vs. 30 mg/kg | *** | 0.0001 |
|  | 3 mg/kg vs. 10 mg/kg | ns | 0.0652 |
|  | 3 mg/kg vs. 30 mg/kg | *** | 0.0005 |
| 60 | Vehicle vs. 1 mg/kg | ns | 0.9657 |
|  | Vehicle vs. 3 mg/kg | ns | 0.8371 |
|  | Vehicle vs. 10 mg/kg | ns | 0.165 |
|  | Vehicle vs. 30 mg/kg | *** | 0.0009 |
|  | 1 mg/kg vs. 3 mg/kg | ns | 0.9412 |
|  | 1 mg/kg vs. 10 mg/kg | ns | 0.1009 |
|  | 1 mg/kg vs. 30 mg/kg | **** | <0.0001 |
|  | 3 mg/kg vs. 10 mg/kg | ns | 0.3917 |
|  | 3 mg/kg vs. 30 mg/kg | *** | 0.0003 |
|  | 10 mg/kg vs. 30 mg/kg | ** | 0.002 |
| **Figure 1F – Male Day 3** | | | |
| **Type of test** | **Comparison** | **P-value summary** | **F, P-value** |
| Two-way ANOVA | Time | **** | F _(3.548, 88.70)_ = 10.15, P<0.0001 |
|  | Treatment | **** | F _(4, 25)_ = 15.83, P=0.1763 |
|  | Interaction | *** | F _(20,125)_ = 10.41, P<0.001 |
| Tukey’s post-hoc |  |  |  |
| 30 | Vehicle vs. 1 mg/kg | ns | 0.0702 |
|  | Vehicle vs. 3 mg/kg | ns | 0.593 |
|  | Vehicle vs. 10 mg/kg | ns | 0.1449 |
|  | Vehicle vs. 30 mg/kg | *** | 0.0009 |
|  | 1 mg/kg vs. 3 mg/kg | ns | >0.9999 |
|  | 1 mg/kg vs. 10 mg/kg | ns | 0.9477 |
|  | 1 mg/kg vs. 30 mg/kg | * | 0.0177 |
|  | 3 mg/kg vs. 10 mg/kg | ns | 0.9898 |
|  | 3 mg/kg vs. 30 mg/kg | ns | 0.0772 |

**Table 2. Statistics from Figure 2**

| **Figure 2D – Male day 1** | | | |
| --- | --- | --- | --- |
| **Type of test** | **Comparison** | **P-value summary** | **F, P-value** |
| Two-way ANOVA | Time | **** | F _(3.282, 82.40)_ = 14.18, P<0.0001 |
|  | Treatment | **** | F _(4, 25)_ = 13.06, P<0.0001 |
|  | Interaction | **** | F _(20,125)_ = 3.948, P<0.0001 |
| Tukey’s post-hoc |  |  |  |
| 30 | Vehicle vs. 1 mg/kg | ns | 0.9015 |
|  | Vehicle vs. 3 mg/kg | ns | 0.0899 |
|  | Vehicle vs. 10 mg/kg | * | 0.0181 |
|  | Vehicle vs. 30 mg/kg | *** | 0.0004 |
|  | 1 mg/kg vs. 3 mg/kg | ns | 0.2789 |
|  | 1 mg/kg vs. 10 mg/kg | * | 0.0448 |
|  | 1 mg/kg vs. 30 mg/kg | ** | 0.001 |
|  | 3 mg/kg vs. 10 mg/kg | ns | 0.4916 |
|  | 3 mg/kg vs. 30 mg/kg | * | 0.0217 |
|  | 10 mg/kg vs. 30 mg/kg | ns | 0.5304 |
| 60 | Vehicle vs. 1 mg/kg | ns | 0.9369 |
|  | Vehicle vs. 3 mg/kg | ns | 0.0732 |
|  | Vehicle vs. 10 mg/kg | ns | 0.1706 |
|  | Vehicle vs. 30 mg/kg | ** | 0.0027 |
|  | 1 mg/kg vs. 3 mg/kg | ns | 0.1214 |
|  | 1 mg/kg vs. 10 mg/kg | ns | 0.1076 |
|  | 1 mg/kg vs. 30 mg/kg | ** | 0.0021 |
|  | 3 mg/kg vs. 10 mg/kg | ns | 0.7871 |
|  | 3 mg/kg vs. 30 mg/kg | * | 0.0286 |
|  | 10 mg/kg vs. 30 mg/kg | ns | 0.3627 |
| 90 | Vehicle vs. 1 mg/kg | ns | 0.9979 |
|  | Vehicle vs. 3 mg/kg | ns | 0.2292 |
|  | Vehicle vs. 10 mg/kg | ns | 0.2059 |
|  | Vehicle vs. 30 mg/kg | ns | 0.0516 |
|  | 1 mg/kg vs. 3 mg/kg | ns | 0.2139 |
|  | 1 mg/kg vs. 10 mg/kg | ns | 0.1909 |
|  | 1 mg/kg vs. 30 mg/kg | * | 0.0452 |
|  | 3 mg/kg vs. 10 mg/kg | ns | >0.9999 |
|  | 3 mg/kg vs. 30 mg/kg | ns | 0.6894 |
|  | 10 mg/kg vs. 30 mg/kg | ns | 0.7628 |
| 180 | Vehicle vs. 1 mg/kg | ns | 0.9504 |
|  | Vehicle vs. 3 mg/kg | ns | 0.9989 |
|  | Vehicle vs. 10 mg/kg | ns | 0.7701 |
|  | Vehicle vs. 30 mg/kg | * | 0.0494 |
|  | 1 mg/kg vs. 3 mg/kg | ns | 0.9922 |
|  | 1 mg/kg vs. 10 mg/kg | ns | 0.9296 |
|  | 1 mg/kg vs. 30 mg/kg | ns | 0.0834 |
|  | 3 mg/kg vs. 10 mg/kg | ns | 0.8474 |
|  | 3 mg/kg vs. 30 mg/kg | ns | 0.0618 |
|  | 10 mg/kg vs. 30 mg/kg | ns | 0.443 |
| **Figure 2E – Female day 1** | | | |
| **Type of test** | **Comparison** | **P-value summary** | **F, P-value** |
| Two-way ANOVA | Time | **** | F _(3.084, 61.67)_ = 12.24, P<0.0001 |
|  | Treatment | **** | F _(4, 20)_ = 7.246, P<0.0001 |
|  | Interaction | **** | F _(20,100)_ = 3.574, P<0.0001 |
| Tukey’s post-hoc |  |  |  |
| 30 | Vehicle vs. 1 mg/kg | ns | 0.9158 |
|  | Vehicle vs. 3 mg/kg | ** | 0.0065 |
|  | Vehicle vs. 10 mg/kg | * | 0.0206 |
|  | Vehicle vs. 30 mg/kg | * | 0.0465 |
|  | 1 mg/kg vs. 3 mg/kg | ns | 0.1389 |
|  | 1 mg/kg vs. 10 mg/kg | ns | 0.1345 |
|  | 1 mg/kg vs. 30 mg/kg | ns | 0.0749 |
|  | 3 mg/kg vs. 10 mg/kg | ns | 0.9949 |
|  | 3 mg/kg vs. 30 mg/kg | ns | 0.4126 |
|  | 10 mg/kg vs. 30 mg/kg | ns | 0.5386 |
| **Figure 2F – Male day 3** | | | |
| **Type of test** | **Comparison** | **P-value summary** | **F, P-value** |
| Two-way ANOVA | Time | ** | F _(3.548, 88.70)_ = 4.806, P=0.0025 |
|  | Treatment | Ns | F _(4, 25)_ = 2.419, P=0.0751 |
|  | Interaction | ns | F _(20,125)_ = 0.9019, P=0.5853 |
| Tukey’s post-hoc |  |  |  |
| 30 | Vehicle vs. 1 mg/kg | ns | 0.9158 |
|  | Vehicle vs. 3 mg/kg | ** | 0.0065 |
|  | Vehicle vs. 10 mg/kg | * | 0.0206 |
|  | Vehicle vs. 30 mg/kg | * | 0.0465 |
|  | 1 mg/kg vs. 3 mg/kg | ns | 0.1389 |
|  | 1 mg/kg vs. 10 mg/kg | ns | 0.1345 |
|  | 1 mg/kg vs. 30 mg/kg | ns | 0.0749 |
|  | 3 mg/kg vs. 10 mg/kg | ns | 0.9949 |
|  | 3 mg/kg vs. 30 mg/kg | ns | 0.4126 |
|  | 10 mg/kg vs. 30 mg/kg | ns | 0.5386 |
| **Figure 2G – Female day 3** | | | |
| **Type of test** | **Comparison** | **P-value summary** | **F, P-value** |
| Two-way ANOVA | Time | * | F _(3.084, 61.67)_ = 3.100, P=0.028 |
|  | Treatment | ns | F _(4, 20)_ = 0.4499, P=0.4499 |
|  | Interaction | ns | F _(20,120)_ = 1.652, P=0.563 |
| Tukey’s post-hoc |  |  |  |
| 30 | Vehicle vs. 1 mg/kg | ns | 0.9158 |
|  | Vehicle vs. 3 mg/kg | ** | 0.0065 |
|  | Vehicle vs. 10 mg/kg | * | 0.0206 |
|  | Vehicle vs. 30 mg/kg | * | 0.0465 |
|  | 1 mg/kg vs. 3 mg/kg | ns | 0.1389 |
|  | 1 mg/kg vs. 10 mg/kg | ns | 0.1345 |
|  | 1 mg/kg vs. 30 mg/kg | ns | 0.0749 |
|  | 3 mg/kg vs. 10 mg/kg | ns | 0.9949 |
|  | 3 mg/kg vs. 30 mg/kg | ns | 0.4126 |
|  | 10 mg/kg vs. 30 mg/kg | ns | 0.5386 |

**Table 3. Statistics from Figure 3**

| **Figure 3B – Male day 1** | | | |
| --- | --- | --- | --- |
| **Type of test** | **Comparison** | **P-value summary** | **F, P-value** |
| Two-way ANOVA | Time | **** | F _(2.902, 46.43)_ = 21.78, P<0.0001 |
|  | Treatment | **** | F _(3, 16)_ = 18.68, P<0.0001 |
|  | Interaction | **** | F _(15,80)_ = 1.582, P<0.0001 |
| Tukey’s post-hoc |  |  |  |
| 30 | Vehicle vs. Δ^9^-THC (3) | ns | 0.9475 |
|  | Vehicle vs. Oxycodone (3) | ** | 0.0038 |
|  | Vehicle vs. Δ^9^-THC (3) + Oxy (3) | *** | 0.0002 |
|  | Δ^9^-THC (3) vs. Oxycodone (3) | ** | 0.0057 |
|  | Δ^9^-THC (3) vs. Δ^9^-THC (3) + Oxy (3) | *** | 0.0003 |
|  | Oxycodone (3) vs. Δ^9^-THC (3) + Oxy (3) | * | 0.0111 |
|  | Vehicle vs. Δ^9^-THC (3) | ** | 0.001 |
| 60 | Vehicle vs. Δ^9^-THC (3) | ns | 0.9523 |
|  | Vehicle vs. Oxycodone (3) | ns | 0.7792 |
|  | Vehicle vs. Δ^9^-THC (3) + Oxy (3) | * | 0.0296 |
|  | Δ^9^-THC (3) vs. Oxycodone (3) | ns | 0.6429 |
|  | Δ^9^-THC (3) vs. Δ^9^-THC (3) + Oxy (3) | * | 0.0195 |
|  | Oxycodone (3) vs. Δ^9^-THC (3) + Oxy (3) | ns | 0.0513 |
| **Figure 3C – Female day 1** | | | |
| **Type of test** | **Comparison** | **P-value summary** | **F, P-value** |
| Two-way ANOVA | Time | **** | F _(2.470, 39.51)_ = 18.72, P<0.0001 |
|  | Treatment | **** | F _(3, 16)_ = 5.458, P<0.0001 |
|  | Interaction | **** | F _(15,80)_ = 5.596, P<0.0001 |
| Tukey’s post-hoc |  |  |  |
| 30 | Vehicle vs. Δ^9^-THC (3) | ns | 0.1566 |
|  | Vehicle vs. Oxycodone (3) | ns | 0.0938 |
|  | Vehicle vs. Δ^9^-THC (3) + Oxy (3) | ** | 0.0016 |
|  | Δ^9^-THC (3) vs. Oxycodone (3) | ns | 0.8679 |
|  | Δ^9^-THC (3) vs. Δ^9^-THC (3) + Oxy (3) | ** | 0.0061 |
|  | Oxycodone (3) vs. Δ^9^-THC (3) + Oxy (3) | * | 0.0108 |
|  | Vehicle vs. Δ^9^-THC (3) | ns | 0.1566 |
| **Figure 3D – Male day 3** | | | |
| **Type of test** | **Comparison** | **P-value summary** | **F, P-value** |
| Two-way ANOVA | Time | ** | F _(2.885, 46.16)_ = 18.72, P=0.0019 |
|  | Treatment | ** | F _(3, 16)_ = 5.605, P=0.0080 |
|  | Interaction | **** | F _(15,80)_ = 4.194, P<0.0001 |
| Tukey’s post-hoc |  |  |  |
| 30 | Vehicle vs. Δ^9^-THC (3) | ns | 0.7055 |
|  | Vehicle vs. Oxycodone (3) | ns | 0.4755 |
|  | Vehicle vs. Δ^9^-THC (3) + Oxy (3) | ** | 0.0025 |
|  | Δ^9^-THC (3) vs. Oxycodone (3) | ns | 0.9616 |
|  | Δ^9^-THC (3) vs. Δ^9^-THC (3) + Oxy (3) | * | 0.011 |
|  | Oxycodone (3) vs. Δ^9^-THC (3) + Oxy (3) | * | 0.0295 |
|  | Vehicle vs. Δ^9^-THC (3) | ns | 0.7055 |
| **Figure 3E – Female day 3** | | | |
| **Type of test** | **Comparison** | **P-value summary** | **F, P-value** |
| Two-way ANOVA | Time | **** | F _(3.393, 54.29)_ = 10.92, P<0.0001 |
|  | Treatment | ns | F _(3, 16)_ = 5.605, P=0.0080 |
|  | Interaction | *** | F _(15,80)_ = 3.268, P<0.001 |
| Tukey’s post-hoc |  |  |  |
| 30 | Vehicle vs. Δ^9^-THC (3) | ns | 0.9999 |
|  | Vehicle vs. Oxycodone (3) | ns | 0.9999 |
|  | Vehicle vs. Δ^9^-THC (3) + Oxy (3) | * | 0.0253 |
|  | Δ^9^-THC (3) vs. Oxycodone (3) | ns | >0.9999 |
|  | Δ^9^-THC (3) vs. Δ^9^-THC (3) + Oxy (3) | ns | 0.0566 |
|  | Oxycodone (3) vs. Δ^9^-THC (3) + Oxy (3) | * | 0.0264 |
|  | Vehicle vs. Δ^9^-THC (3) | ns | 0.9999 |

**Table 4. Statistics from Figure 4**

| **Figure 4B – Oxycodone Withdrawal Males** | | | |
| --- | --- | --- | --- |
| **Type of test** | **Comparison** | **P-value summary** | **F, P-value** |
| One-way ANOVA | All groups | **** | F _(4, 24)_ = 5.039, P=0.0043 |
| Tukey’s post-hoc | 0 vs. 1 | ns | 0.2972 |
|  | 0 vs. 3 | ns | 0.0577 |
|  | 0 vs. 10 | * | 0.0389 |
|  | 0 vs. 30 | ** | 0.0024 |
|  | 1 vs. 3 | ns | 0.9009 |
|  | 1 vs. 10 | ns | 0.8225 |
|  | 1 vs. 30 | ns | 0.1687 |
|  | 3 vs. 10 | ns | 0.9997 |
|  | 3 vs. 30 | ns | 0.5759 |
|  | 10 vs. 30 | ns | 0.6821 |
| **Figure 4C – Oxycodone Withdrawal Females** | | | |
| **Type of test** | **Comparison** | **P-value summary** | **F, P-value** |
| One-way ANOVA | All groups | **** | F _(4, 20)_ = 4.692, P=0.0078 |
| Tukey’s post-hoc | 0 vs. 1 | ns | 0.7539 |
|  | 0 vs. 3 | ns | 0.724 |
|  | 0 vs. 10 | * | 0.0447 |
|  | 0 vs. 30 | * | 0.0243 |
|  | 1 vs. 3 | ns | >0.9999 |
|  | 1 vs. 10 | ns | 0.312 |
|  | 1 vs. 30 | ns | 0.1904 |
|  | 3 vs. 10 | ns | 0.3387 |
|  | 3 vs. 30 | ns | 0.2093 |
|  | 10 vs. 30 | ns | 0.9978 |
| **Figure 4D – Combo Withdrawal Males** | | | |
| **Type of test** | **Comparison** | **P-value summary** | **F, P-value** |
| One-way ANOVA | All groups | ** | F _(3, 16)_ = 7.853, P=0.0019 |
| Tukey’s post-hoc | Vehicle vs. Δ^9^-THC (3) | ns | >0.9999 |
|  | Vehicle vs. Oxy (3) | ** | 0.0034 |
|  | Vehicle vs. Oxy (3) +THC (3) | ns | 0.5582 |
|  | Δ^9^-THC (3) vs. Oxy (3) | ** | 0.0034 |
|  | Δ^9^-THC (3) vs. Oxy (3) +THC (3) | ns | 0.5582 |
|  | Oxy(3) vs. Oxy(3) +THC (3) | * | 0.0489 |
| **Figure 4D – Combo Withdrawal Males** | | | |
| **Type of test** | **Comparison** | **P-value summary** | **F, P-value** |
| One-way ANOVA | All groups | ** | F _(3, 16)_ = 5.300, P=0.0099 |
| Tukey’s post-hoc | Vehicle vs. Δ^9^-THC (3) | ns | 0.999 |
|  | Vehicle vs. Oxy (3) | ns | 0.0794 |
|  | Vehicle vs. Oxy (3) +THC (3) | * | 0.0493 |
|  | Δ^9^-THC (3) vs. Oxy (3) | ns | 0.0611 |
|  | Δ^9^-THC (3) vs. Oxy (3) +THC (3) | * | 0.0376 |
|  | Oxy(3) vs. Oxy(3) +THC (3) | ns | 0.9941 |

**Table 5. Statistics from Figure 5**

| **Figure 5G – Circadian Activity Index Males** | | | |
| --- | --- | --- | --- |
| **Type of test** | **Comparison** | **P-value summary** | **F, P-value** |
| Two-way ANOVA | Time | **** | F _(2.568, 41.09)_ = 87.21, P<0.0001 |
|  | Treatment | **** | F _(3, 16)_ = 15.52, P<0.0001 |
|  | Interaction | **** | F _(9,48)_ = 7.326, P<0.0001 |
| Tukey’s post-hoc |  |  |  |
| Baseline | Vehicle vs. Δ^9^-THC (3) | ns | 0.639 |
|  | Vehicle vs. Oxycodone (3) | ns | 0.8054 |
|  | Vehicle vs. Δ^9^-THC (3) + Oxy (3) | ns | 0.9986 |
|  | Δ^9^-THC (3) vs. Oxycodone (3) | ns | 0.9975 |
|  | Δ^9^-THC (3) vs. Δ^9^-THC (3) + Oxy (3) | ns | 0.4463 |
|  | Oxycodone (3) vs. Δ^9^-THC (3) + Oxy (3) | ns | 0.7425 |
| Dosing | Vehicle vs. Δ9-THC (3) | ns | 0.0747 |
|  | Vehicle vs. Oxycodone (3) | ** | 0.0018 |
|  | Vehicle vs. Δ9-THC (3) + Oxy (3) | **** | <0.0001 |
|  | Δ9-THC (3) vs. Oxycodone (3) | * | 0.0127 |
|  | Δ9-THC (3) vs. Δ9-THC (3) + Oxy (3) | *** | 0.0008 |
|  | Oxycodone (3) vs. Δ9-THC (3) + Oxy (3) | ns | 0.5293 |
| Withdrawal | Vehicle vs. Δ9-THC (3) | ns | 0.9893 |
|  | Vehicle vs. Oxycodone (3) | ns | 0.0771 |
|  | Vehicle vs. Δ9-THC (3) + Oxy (3) | ** | 0.0093 |
|  | Δ9-THC (3) vs. Oxycodone (3) | ns | 0.1437 |
|  | Δ9-THC (3) vs. Δ9-THC (3) + Oxy (3) | * | 0.036 |
|  | Oxycodone (3) vs. Δ9-THC (3) + Oxy (3) | ns | 0.9666 |
| Recovery | Vehicle vs. Δ9-THC (3) | ns | 0.9968 |
|  | Vehicle vs. Oxycodone (3) | ns | 0.3531 |
|  | Vehicle vs. Δ9-THC (3) + Oxy (3) | ns | 0.995 |
|  | Δ9-THC (3) vs. Oxycodone (3) | ns | 0.397 |
|  | Δ9-THC (3) vs. Δ9-THC (3) + Oxy (3) | ns | 0.9762 |
|  | Oxycodone (3) vs. Δ9-THC (3) + Oxy (3) | ns | 0.391 |
| **Figure 5H – Circadian Activity Index Females** | | | |
| **Type of test** | **Comparison** | **P-value summary** | **F, P-value** |
| Two-way ANOVA | Time | **** | F _(2.505, 40.08)_ = 31.95, P<0.0001 |
|  | Treatment | ns | F _(3, 16)_ = 1.591, P=0.2307 |
|  | Interaction | * | F _(9,48)_ = 2.360, P=0.0269 |
| Tukey’s post-hoc |  |  |  |
| Baseline | Vehicle vs. Δ^9^-THC (3) | ns | 0.9885 |
|  | Vehicle vs. Oxycodone (3) | ns | 0.9877 |
|  | Vehicle vs. Δ^9^-THC (3) + Oxy (3) | ns | 0.9674 |
|  | Δ^9^-THC (3) vs. Oxycodone (3) | ns | >0.9999 |
|  | Δ^9^-THC (3) vs. Δ^9^-THC (3) + Oxy (3) | ns | 0.9973 |
|  | Oxycodone (3) vs. Δ^9^-THC (3) + Oxy (3) | ns | 0.9908 |
| Dosing | Vehicle vs. Δ9-THC (3) | ns | 0.6514 |
|  | Vehicle vs. Oxycodone (3) | * | 0.0275 |
|  | Vehicle vs. Δ9-THC (3) + Oxy (3) | * | 0.0151 |
|  | Δ9-THC (3) vs. Oxycodone (3) | ns | 0.0524 |
|  | Δ9-THC (3) vs. Δ9-THC (3) + Oxy (3) | ** | 0.0077 |
|  | Oxycodone (3) vs. Δ9-THC (3) + Oxy (3) | ns | 0.7014 |
| Withdrawal | Vehicle vs. Δ9-THC (3) | ns | 0.9984 |
|  | Vehicle vs. Oxycodone (3) | ns | 0.9376 |
|  | Vehicle vs. Δ9-THC (3) + Oxy (3) | ns | 0.9126 |
|  | Δ9-THC (3) vs. Oxycodone (3) | ns | 0.4898 |
|  | Δ9-THC (3) vs. Δ9-THC (3) + Oxy (3) | ns | 0.0763 |
|  | Oxycodone (3) vs. Δ9-THC (3) + Oxy (3) | ns | 0.9666 |
| Recovery | Vehicle vs. Δ9-THC (3) | ns | >0.9999 |
|  | Vehicle vs. Oxycodone (3) | ns | >0.9999 |
|  | Vehicle vs. Δ9-THC (3) + Oxy (3) | ns | 0.7981 |
|  | Δ9-THC (3) vs. Oxycodone (3) | ns | >0.9999 |
|  | Δ9-THC (3) vs. Δ9-THC (3) + Oxy (3) | ns | 0.6732 |
|  | Oxycodone (3) vs. Δ9-THC (3) + Oxy (3) | ns | 0.6213 |

**Table 6. Statistics from Figure 6**

| **Figure 6C – Male Oxycodone (3)** | | | |
| --- | --- | --- | --- |
| **Type of test** | **Comparison** | **P-value summary** | **F, P-value** |
| Two-way ANOVA | Time | ns | F _(1,12)_ = 0.0264, P=0.8736 |
|  | Treatment | ns | F _(1,12)_ = 2.435, P=0.1446 |
|  | Interaction | **** | F _(1,12)_ = 48.94, P<0.0001 |
| Sidak post-hoc |  |  |  |
| Pre-test | Vehicle vs. Oxycodone (3) | ns | 0.3803 |
| Post-test | Vehicle vs. Oxycodone (3) | *** | 0.0007 |
| **Figure 6D – Female Oxycodone (3)** | | | |
| **Type of test** | **Comparison** | **P-value summary** | **F, P-value** |
| Two-way ANOVA | Time | ns | F _(1,14)_ = 0.01402, P=0.9074 |
|  | Treatment | ns | F _(1,14)_ = 3.140, P=0.0982 |
|  | Interaction | **** | F _(1,14)_ = 81.60, P<0.0001 |
| Sidak post-hoc |  |  |  |
| Pre-test | Vehicle vs. Oxycodone (3) | ns | 0.2123 |
| Post-test | Vehicle vs. Oxycodone (3) | **** | <0.0001 |
| **Figure 6E –Male Δ^9^-THC (3) + Oxycodone (1)** | | | |
| **Type of test** | **Comparison** | **P-value summary** | **F, P-value** |
| Two-way ANOVA | Time | ns | F _(1,12)_ = 0.0023, P=0.8831 |
|  | Treatment | ns | F _(1,12)_ = 2.678, P=0.1277 |
|  | Interaction | * | F _(1,12)_ = 5.241, P=0.0421 |
| Sidak post-hoc |  |  |  |
| Pre-test | Vehicle vs. Δ9-THC (3) + Oxycodone (1) | ns | 0.9658 |
| Post-test | Vehicle vs. Δ9-THC (3) + Oxycodone (1) | * | 0.0328 |
| **Figure 6F – Female Δ^9^-THC (3) + Oxycodone (1)** | | | |
| **Type of test** | **Comparison** | **P-value summary** | **F, P-value** |
| Two-way ANOVA | Time | ns | F _(1,10)_ = 0.005, P=0.9427 |
|  | Treatment | ns | F _(1,10)_ = 2.501, P=0.1449 |
|  | Interaction | **** | F _(1,10)_ = 6.729, P=0.0268 |
| Sidak post-hoc |  |  |  |
| Pre-test | Vehicle vs. Δ9-THC (3) + Oxycodone (1) | ns | 0.9836 |
| Post-test | Vehicle vs. Δ9-THC (3) + Oxycodone (1) | * | 0.0234 |
| **Figure 6G – Male Δ^9^-THC (3) + Oxycodone (3)** | | | |
| **Type of test** | **Comparison** | **P-value summary** | **F, P-value** |
| Two-way ANOVA | Time | ns | F _(1,14)_ = 0.003, P=0.8572 |
|  | Treatment | ** | F _(1,14)_ = 11.36, P=0.0046 |
|  | Interaction | **** | F _(1,14)_ = 41.26, P<0.0001 |
| Sidak post-hoc |  |  |  |
| Pre-test | Vehicle vs. Δ9-THC (3) + Oxycodone (3) | ns | 0.9858 |
| Post-test | Vehicle vs. Δ9-THC (3) + Oxycodone (3) | **** | <0.0001 |
| **Figure 6H –Female Δ^9^-THC (3) + Oxycodone (3)** | | | |
| **Type of test** | **Comparison** | **P-value summary** | **F, P-value** |
| Two-way ANOVA | Time | ns | F _(1,14)_ = 0.1671, P=0.6889 |
|  | Treatment | ns | F _(1,14)_ = 0.8393, P=0.3751 |
|  | Interaction | * | F _(1,14)_ = 8.366, P=0.0118 |
| Sidak post-hoc |  |  |  |
| Pre-test | Vehicle vs. Δ9-THC (3) + Oxycodone (3) | ns | 0.5933 |
| Post-test | Vehicle vs. Δ9-THC (3) + Oxycodone (3) | * | 0.044 |

**Table 7. Statistics from Figure 7**

| **Figure 7A – Locomotor Sensitization Males** | | | |
| --- | --- | --- | --- |
| **Type of test** | **Comparison** | **P-value summary** | **F, P-value** |
| Two-way ANOVA | Time | * | F _(1.578, 61.54)_ = 4.927, P=0.0161 |
|  | Treatment | **** | F _(5, 39)_ = 43.00, P<0.0001 |
|  | Interaction | **** | F _(10,78)_ = 4.995, P<0.0001 |
| Tukey’s post-hoc |  |  |  |
| Day 1 | Vehicle vs. Oxycodone (1) | ns | 0.972 |
|  | Vehicle vs. Oxycodone (3) | ** | 0.0075 |
|  | Vehicle vs. Δ^9^-THC (3) | ns | 0.1219 |
|  | Vehicle vs. Oxycodone (1)+Δ^9^-THC (3) | ns | 0.7807 |
|  | Vehicle vs. Oxycodone (3)+Δ^9^-THC (3) | * | 0.0166 |
|  | Oxycodone (1) vs. Oxycodone (3) | * | 0.0103 |
|  | Oxycodone (1) vs. Δ^9^-THC (3) | * | 0.0459 |
|  | Oxycodone (1) vs. Oxycodone (1)+Δ^9^-THC (3) | ns | 0.9636 |
|  | Oxycodone (1) vs. Oxycodone (3)+Δ^9^-THC (3) | * | 0.0265 |
|  | Oxycodone (3) vs. Δ^9^-THC (3) | ** | 0.0031 |
|  | Oxycodone (3) vs. Oxycodone (1)+Δ^9^-THC (3) | * | 0.0202 |
|  | Oxycodone (3) vs. Oxycodone (3)+Δ^9^-THC (3) | ns | 0.7934 |
|  | Δ^9^-THC (3) vs. Oxycodone (1)+Δ^9^-THC (3) | ns | 0.1286 |
|  | Δ^9^-THC (3) vs. Oxycodone (3)+Δ^9^-THC (3) | ** | 0.0047 |
|  | Oxycodone (1)+Δ^9^-THC (3) vs. Oxycodone (3)+Δ^9^-THC (3) | ns | 0.0862 |
| Day 2 | Vehicle vs. Oxycodone (1) | ns | 0.9372 |
|  | Vehicle vs. Oxycodone (3) | ** | 0.0016 |
|  | Vehicle vs. Δ9-THC (3) | ns | 0.2269 |
|  | Vehicle vs. Oxycodone (1)+Δ9-THC (3) | ns | 0.4254 |
|  | Vehicle vs. Oxycodone (3)+Δ9-THC (3) | *** | 0.0002 |
|  | Oxycodone (1) vs. Oxycodone (3) | ** | 0.002 |
|  | Oxycodone (1) vs. Δ9-THC (3) | ns | 0.1176 |
|  | Oxycodone (1) vs. Oxycodone (1)+Δ9-THC (3) | ns | 0.7444 |
|  | Oxycodone (1) vs. Oxycodone (3)+Δ9-THC (3) | *** | 0.0002 |
|  | Oxycodone (3) vs. Δ9-THC (3) | ** | 0.001 |
|  | Oxycodone (3) vs. Oxycodone (1)+Δ9-THC (3) | ** | 0.0044 |
|  | Oxycodone (3) vs. Oxycodone (3)+Δ9-THC (3) | ns | 0.9988 |
|  | Δ9-THC (3) vs. Oxycodone (1)+Δ9-THC (3) | ns | 0.11 |
|  | Δ9-THC (3) vs. Oxycodone (3)+Δ9-THC (3) | *** | 0.0001 |
|  | Oxycodone (1)+Δ9-THC (3) vs. Oxycodone (3)+Δ9-THC (3) | *** | 0.0008 |
| Day 3 | Vehicle vs. Oxycodone (1) | ns | >0.9999 |
|  | Vehicle vs. Oxycodone (3) | *** | 0.0003 |
|  | Vehicle vs. Δ9-THC (3) | * | 0.0399 |
|  | Vehicle vs. Oxycodone (1)+Δ9-THC (3) | ns | 0.9992 |
|  | Vehicle vs. Oxycodone (3)+Δ9-THC (3) | ** | 0.0083 |
|  | Oxycodone (1) vs. Oxycodone (3) | *** | 0.0005 |
|  | Oxycodone (1) vs. Δ9-THC (3) | **** | <0.0001 |
|  | Oxycodone (1) vs. Oxycodone (1)+Δ9-THC (3) | ns | 0.9997 |
|  | Oxycodone (1) vs. Oxycodone (3)+Δ9-THC (3) | ** | 0.0089 |
|  | Oxycodone (3) vs. Δ9-THC (3) | *** | 0.0002 |
|  | Oxycodone (3) vs. Oxycodone (1)+Δ9-THC (3) | *** | 0.0002 |
|  | Oxycodone (3) vs. Oxycodone (3)+Δ9-THC (3) | ns | 0.955 |
|  | Δ9-THC (3) vs. Oxycodone (1)+Δ9-THC (3) | ns | 0.0532 |
|  | Δ9-THC (3) vs. Oxycodone (3)+Δ9-THC (3) | ** | 0.0026 |
|  | Oxycodone (1)+Δ9-THC (3) vs. Oxycodone (3)+Δ9-THC (3) | ** | 0.007 |
| **Figure 7B – Locomotor Sensitization Females** | | | |
| **Type of test** | **Comparison** | **P-value summary** | **F, P-value** |
| Two-way ANOVA | Time | ns | F _(1.944, 56.38)_ = 3.188, P=0.0501 |
|  | Treatment | **** | F _(5, 39)_ = 12.29, P<0.0001 |
|  | Interaction | ns | F _(10,58)_ = 0.9856, P=0.4661 |
| Tukey’s post-hoc |  |  |  |
| Day 1 | Vehicle vs. Oxycodone (1) | ns | 0.3059 |
|  | Vehicle vs. Oxycodone (3) | *** | 0.0003 |
|  | Vehicle vs. Δ^9^-THC (3) | ns | 0.6909 |
|  | Vehicle vs. Oxycodone (1)+Δ^9^-THC (3) | ns | 0.9933 |
|  | Vehicle vs. Oxycodone (3)+Δ^9^-THC (3) | ** | 0.0028 |
|  | Oxycodone (1) vs. Oxycodone (3) | * | 0.044 |
|  | Oxycodone (1) vs. Δ^9^-THC (3) | ns | 0.1359 |
|  | Oxycodone (1) vs. Oxycodone (1)+Δ^9^-THC (3) | ns | 0.6764 |
|  | Oxycodone (1) vs. Oxycodone (3)+Δ^9^-THC (3) | * | 0.0373 |
|  | Oxycodone (3) vs. Δ^9^-THC (3) | *** | 0.0004 |
|  | Oxycodone (3) vs. Oxycodone (1)+Δ^9^-THC (3) | ** | 0.002 |
|  | Oxycodone (3) vs. Oxycodone (3)+Δ^9^-THC (3) | ns | 0.9642 |
|  | Δ^9^-THC (3) vs. Oxycodone (1)+Δ^9^-THC (3) | ns | 0.7059 |
|  | Δ^9^-THC (3) vs. Oxycodone (3)+Δ^9^-THC (3) | *** | 0.001 |
|  | Oxycodone (1)+Δ^9^-THC (3) vs. Oxycodone (3)+Δ^9^-THC (3) | ** | 0.004 |
| Day 2 | Vehicle vs. Oxycodone (1) | ns | 0.1238 |
|  | Vehicle vs. Oxycodone (3) | * | 0.0268 |
|  | Vehicle vs. Δ9-THC (3) | ns | 0.9963 |
|  | Vehicle vs. Oxycodone (1)+Δ9-THC (3) | ns | 0.5619 |
|  | Vehicle vs. Oxycodone (3)+Δ9-THC (3) | ** | 0.0016 |
|  | Oxycodone (1) vs. Oxycodone (3) | ns | 0.1583 |
|  | Oxycodone (1) vs. Δ9-THC (3) | ns | 0.0823 |
|  | Oxycodone (1) vs. Oxycodone (1)+Δ9-THC (3) | ns | 0.9715 |
|  | Oxycodone (1) vs. Oxycodone (3)+Δ9-THC (3) | * | 0.011 |
|  | Oxycodone (3) vs. Δ9-THC (3) | * | 0.0241 |
|  | Oxycodone (3) vs. Oxycodone (1)+Δ9-THC (3) | ns | 0.0963 |
|  | Oxycodone (3) vs. Oxycodone (3)+Δ9-THC (3) | ns | 0.9677 |
|  | Δ9-THC (3) vs. Oxycodone (1)+Δ9-THC (3) | ns | 0.397 |
|  | Δ9-THC (3) vs. Oxycodone (3)+Δ9-THC (3) | ** | 0.0018 |
|  | Oxycodone (1)+Δ9-THC (3) vs. Oxycodone (3)+Δ9-THC (3) | ** | 0.0058 |
| Day 3 | Vehicle vs. Oxycodone (1) | ns | 0.1898 |
|  | Vehicle vs. Oxycodone (3) | * | 0.0131 |
|  | Vehicle vs. Δ9-THC (3) | ns | 0.46 |
|  | Vehicle vs. Oxycodone (1)+Δ9-THC (3) | ns | 0.5221 |
|  | Vehicle vs. Oxycodone (3)+Δ9-THC (3) | * | 0.0315 |
|  | Oxycodone (1) vs. Oxycodone (3) | ns | 0.1528 |
|  | Oxycodone (1) vs. Δ9-THC (3) | * | 0.0332 |
|  | Oxycodone (1) vs. Oxycodone (1)+Δ9-THC (3) | ns | 0.9606 |
|  | Oxycodone (1) vs. Oxycodone (3)+Δ9-THC (3) | ns | 0.1223 |
|  | Oxycodone (3) vs. Δ9-THC (3) | ** | 0.0053 |
|  | Oxycodone (3) vs. Oxycodone (1)+Δ9-THC (3) | ns | 0.069 |
|  | Oxycodone (3) vs. Oxycodone (3)+Δ9-THC (3) | ns | 0.8586 |
|  | Δ9-THC (3) vs. Oxycodone (1)+Δ9-THC (3) | ns | 0.0654 |
|  | Δ9-THC (3) vs. Oxycodone (3)+Δ9-THC (3) | * | 0.0178 |
|  | Oxycodone (1)+Δ9-THC (3) vs. Oxycodone (3)+Δ9-THC (3) | ns | 0.0779 |
